# Clogging of particle suspensions in networks

**DOI:** 10.64898/2026.09.07.749836

**Authors:** Cara V. Neal, Duncan R. Hewitt, Philip Pearce

## Abstract

In many biological, biomedical and industrial systems, particles are transported via fluids through confined networks, in which clogging can disrupt function. However, we lack a predictive theoretical framework that couples particle transport, suspension rheology and network flow resistance. Here, we develop a model and solution algorithm for particle suspension flow in networks based on vessel-level continuum modelling and particle distribution at nodes connecting vessels. We apply the model to study transport of dense particle suspensions in minimal and physiological biological networks. A key feature of our model is the coupling between particle volume fraction and particle flux: in line with the physics of dense suspensions, each network branch, or vessel, possesses a local carrying capacity for particle transport at an intermediate particle fraction between zero and the maximum packing fraction. If this flux capacity is reached, the vessel becomes flux-limited and particles can accumulate in upstream branches, causing them to enter a high-particle-fraction, high-resistance state that we refer to as ‘clogged’. We show that these vessel flux limitations lead to network-level redistribution of particles, which can cause widespread clogging and emergent network-scale heterogeneity. By varying network topology, we find that in some regimes increasing network connectivity does not improve transport: paradoxically, additional pathways can promote clogging and reduce network-level particle flux, analogous to classic results in traffic flow networks. Our results provide a minimal mechanistic framework that links suspension physics, network topology, and transport failure in complex flow networks.

## I. INTRODUCTION

Many natural and engineered systems transport particles via fluid flow through branching networks. Examples include filtration systems [1], microfluidic devices [2], animal circulatory systems [3, 4], plant vasculature [5–7], and fungal transport networks [8–11]. Transport in these systems is governed by both suspension dynamics and network architecture. Studies of flow through transport networks have shown that network structure controls efficiency, robustness, and flow distribution, with, for example, loops and redundant pathways playing important roles in resilience to damage and in containing the effects of local blockages [12–18]. Under confinement, particle suspensions can exhibit accumulation, bridging, and aggregation due to hydrodynamic interactions, shear-induced migration, and inter-particle forces [19–21]. These processes can lead to ‘clogging’: the formation of an obstruction, typically at a constriction, which sharply increases local resistance, suppresses downstream transport, and promotes upstream particle accumulation [22] (see Fig. 1a). In applications such as filtration and microfluidic cell sorting, clogging can be exploited to contribute to network function [23, 24], but it more commonly reduces transport efficiency and contributes to dysfunction in biological and porous transport systems [25–28]. Although the influence of network architecture on transport efficiency and robustness is well established in Newtonian flow networks [14, 29], how suspension-driven transport limitations interact with network structure remains poorly understood.

**FIG. 1.**
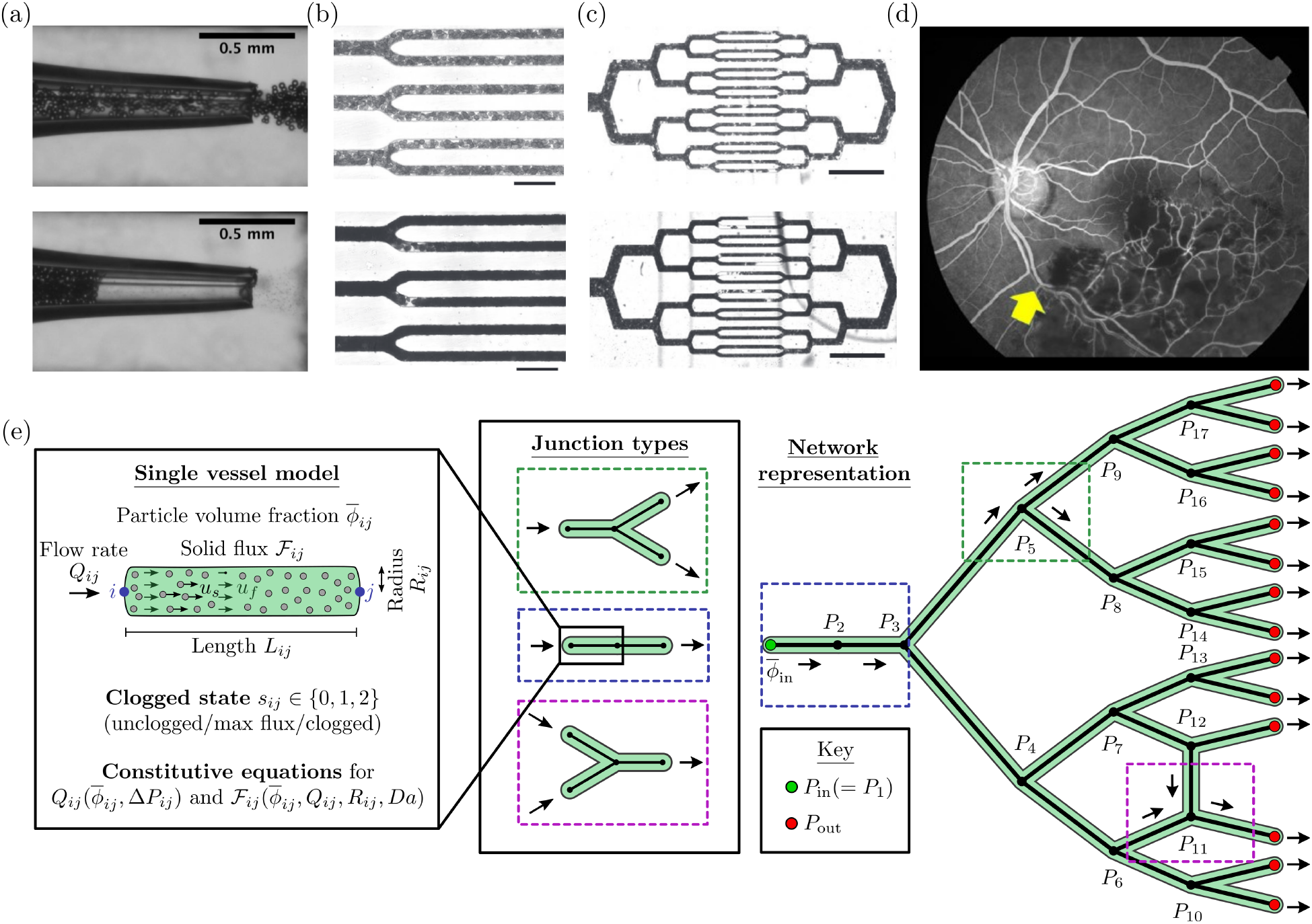
Multiscale clogging in biological transport networks. (a) Clogging via particle bridging in a tapered glass capillary, illustrating flow disruption in confined geometries. Top: freely flowing suspension. Bottom: particle bridging at the constriction, forming a clog that obstructs down-stream transport (reproduced, with permission, from [19]). (b–c) Flow and occlusion of sickle-cell blood in a microfluidic network representing the microvasculature under varying oxygen tension (high: top; low: bottom) (reproduced, with permission, from [30]). Panel (b) shows junction-level occlusion at higher magnification, while (c) shows network-scale disruption. Scale bars: 20*µ*m in (b) and 200*µ*m in (c). (d) Retinal vasculature occlusion demonstrating downstream perfusion loss (reproduced from [31] under the Creative Commons CC BY 4.0 licence). Dark regions show reduced perfusion, occurring predominantly downstream of the occluded vessel marked by the yellow arrow, but also extending into surrounding tissue. (e) Schematic of the modelling framework here. Single vessel flow between nodes *i* and *j* is described by constitutive equations for *Q*_*ij*_ and ℱ_*ij*_, with a maximum solid flux 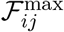 and vessel clogged state *s*_*ij*_. Junctions obey conservation laws coupled with flux constraints, and are assembled into network-scale models of transport and occlusion.

The interplay between clogging and transport through networks is evident across a wide range of biological systems. In bacterial systems, biofilms and filamentous biofilm streamers can trigger sudden clogging and flow distribution in porous media and microfluidic devices [32, 33]. In filamentous fungi, septal pores connecting neighbouring cellular compartments can be rapidly sealed by particles called Woronin bodies following damage [34]. In the microvasculature, vessels can be occluded by the blood, which is a dense suspension of red blood cells (RBC) and other constituents such as plasma. Vessel occlusion, or vaso-occlusion, is promoted in diseases that affect RBC deformability, such as diabetes [35] and sickle cell disease, a hereditary condition in which deoxygenation causes haemoglobin polymerisation and RBC stiffening [36]. Increased adhesion to vessel walls and altered blood rheology further enhance the likelihood of vaso-occlusion in sickle cell disease [37]. Microfluidic studies have shown how vascular architecture and oxygen tension regulate sickle-cell blood flow and clogging [30, 38], revealing both local junction-level occlusion and large-scale disruption of network flow, with clogs observed forming upstream of capillaries in low oxygen tension blood (Fig. 1b–c). More widely, in retinal networks [31], pathologies such as diabetic retinopathy and retinal vein occlusion [39, 40] can involve vaso-occlusion causing reduced perfusion downstream and in neighbouring tissue (Fig. 1d); however, we lack a clear mechanistic basis for these pathologies, to identify new risk factors and potential clinical interventions [41]. Together, these studies motivate the need for predictive models that couple suspension rheology and clogging with network-scale flow redistribution.

Stochastic and discrete methods have been developed to study suspension flow through constricted geometries [42–47], but are computationally expensive for network-scale problems. Continuum approaches offer an alternative route for modelling dense suspensions [48]. In particular, Herale *et al*. [49] developed a minimal two-phase model for suspension flow in a channel with a constriction, consisting of a particle-laden ‘wet solid’ phase coupled to a Darcy seepage flow through the moving solid particles [49]. A feature of this model is that the solid flux is non-monotonic in particle volume fraction, leading to a maximum solid flux that the vessel can sustain. Beyond this limit, additional particles cannot be accommodated downstream of the constriction and instead accumulate upstream in a high-volume-fraction, high-resistance state referred to as a clog. The emergence of such flux-limited states is qualitatively consistent with experimentally observed transitions in dense suspension flows under confinement [22]. In terms of networks, we expect inherently non-local effects: a blockage in one branch will redistribute flux, potentially triggering secondary blockages. For example, experimental studies of colloidal suspension flow through parallel microchannels show that clog growth in one channel can redistribute flow to neighbouring channels [50]. In recent work, clogging by individual cells has been incorporated into stochastic models of microfluidic networks [51]; however, we lack predictive frameworks that couple suspension rheology, flux redistribution, and clogging in general branching networks with larger vessels.

In this work, we develop a continuum framework for dense-suspension transport in branching networks that couples vessel-scale suspension rheology to network-scale flow and particle redistribution (Fig. 1e). Building on the single-vessel model of Herale *et al*. [49], we introduce the concept of a vessel-scale maximum solid flux into a network setting, allowing local transport limitations to generate upstream accumulation and clogging. Using simple network motifs, we show how local flux constraints generate both local and non-local redistribution of particles, leading to clogging and large-scale changes in transport behaviour. We further demonstrate that increasing network connectivity does not necessarily improve transport, with additional pathways either alleviating or promoting clogging depending on how they redistribute flux relative to downstream transport capacities. Finally, we apply the frame-work to a bio-inspired vascular network, to predict how local flux limitations can combine to generate large-scale transport disruption through non-local clogging.

## II. MODEL FORMULATION

### A. Overview of model

We model dense particle suspension transport through a branching network using a one-dimensional continuum approach that captures network-level distribution of solid flux and the onset of clogging without explicitly resolving particle-scale dynamics (Fig. 1e). The network is represented as directed and acyclic with a single inlet, in which edges *V*_*ij*_ represent vessels connecting pressure nodes *i* and *j*. Here, the term ‘vessel’ is used as a generic term for a network segment and could represent a blood vessel, microfluidic channel or other network transport element depending on the application. Each vessel has prescribed geometry (length *L*_*ij*_, radius *R*_*ij*_) and is associated with an unknown steady total volumetric flow rate *Q*_*ij*_, solid flux ℱ_*ij*_, and pipe-averaged particle volume fraction 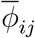. Pressures *P*_*i*_ are defined at nodes, and flow in each vessel is driven by the pressure difference between nodes, Δ*P*_*ij*_. The total pressure drop across the network is labelled Δ*P*_total_ = *P*_in_ − *P*_out_, where *P*_in_(= *P*_1_) and *P*_out_ are the inlet and outlet pressures. The inlet particle volume fraction is prescribed as 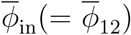, although we will later find that particle accumulation can increase the inlet particle volume fraction above this value.

A steady network state is obtained by combining several key ingredients: vessel-scale constitutive laws that relate the pressure drop and particle volume fraction 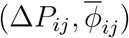 to flow rate and particle flux (*Q*_*ij*_, ℱ_*ij*_); capacity constraints on per-vessel solid flux; junction conservation of mass for *Q*_*ij*_ and ℱ_*ij*_; a flux splitting rule at nodes; and a branch selection rule that represents clogging within the continuum framework.

A key feature of the vessel constitutive laws is the existence of a finite maximum solid flux, 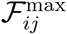, which depends on both vessel geometry and suspension properties and defines a local carrying capacity for particle transport. This flux-limiting behaviour originates from the physics of dense suspensions, as modelled by Herale *et al*. [49], in which transport limitations arise due to a geometric constriction within an isolated vessel. Here, instead of modelling local constrictions within individual vessels, we treat each vessel as a uniform segment characterised by its geometry and investigate how network heterogeneity in vessel capacities drives redistribution of solid flux across the network, leading to non-local effects on transport. At junctions, incoming solid flux is partitioned among outgoing vessels according to a flux-splitting law which is adjusted, if necessary, to satisfy downstream capacity constraints. If the incoming solid flux exceeds the total downstream carrying capacity, particles cannot be transmitted further downstream and must accumulate upstream. Within the present framework, this accumulation is represented through a transition from a 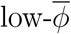 to a 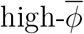 solution branch. We refer to vessels occupying this 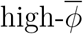 state as ‘clogged’. In this terminology, clogging does not imply complete blockage of transport, but rather a state of particle accumulation associated with high resistance. Together, these ingredients define a non-linear network problem for the flow, solid flux, and particle volume fraction.

### B. Vessel-scale transport and flux capacity constraints

To calculate flow and transport in each vessel, we use constitutive relations derived and non-dimensionalised in Appendix A, which are based on a simplified version of the dense-suspension model of Herale *et al*. [49]. Unless otherwise stated, all quantities in the remainder of the manuscript are dimensionless and scaled with respect to the inlet vessel. In the model, the motion of the suspension is described by two parallel transport pathways: a particle-rich ‘wet solid’ phase and a Darcy seepage flow of interstitial fluid through the suspension. Consequently, each vessel *V*_*ij*_ is characterised by a suspension resistance 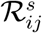 and a Darcy resistance 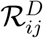, which act in parallel to determine the total flow rate. The resulting constitutive relations give the total volumetric flow rate *Q*_*ij*_ and solid flux ℱ_*ij*_ in terms of the pressure drop Δ*P*_*ij*_, vessel geometry, and particle volume fraction 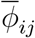:

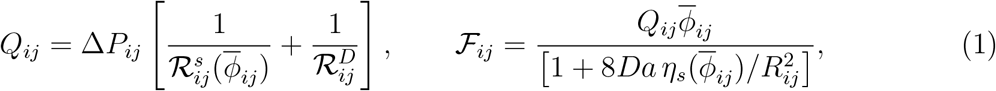

where

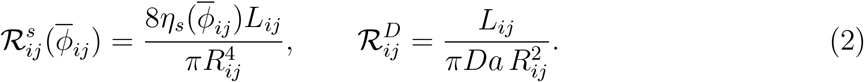

The suspension resistance depends on the particle volume fraction through the effective suspension viscosity (*η*_*s*_), which is modelled as

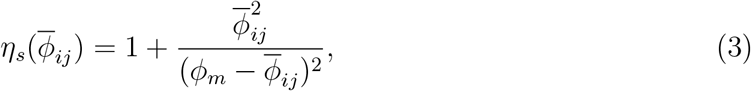

(see Appendix A), where *ϕ*_*m*_ is the maximum packing fraction of particles. The Darcy resistance depends on the Darcy number 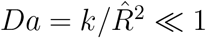, which determines the influence of permeability on suspension transport, where *k* is the permeability (assumed constant here) and 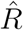 is radius of the inlet vessel.

For fixed (*Q, R, Da*), the flux 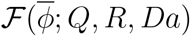 is non-monotonic as the particle volume fraction 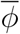 is varied (see, for example, Fig. 2). When 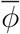 is not near *ϕ*_*m*_, the suspension resistance is small relative to the Darcy resistance, and ℱ increases with 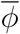. As 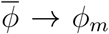, however, the rapid growth of 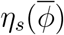 leads to a sharp increase in suspension resistance, reducing the suspension velocity. In this limit, the total flux is dominated by Darcy flow and ℱ is reduced (ultimately to zero as 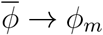). We therefore define the per-vessel solid flux capacity

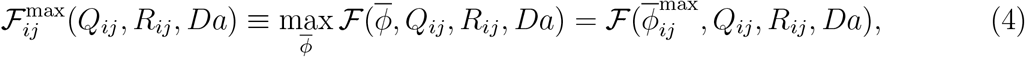

where 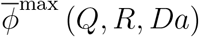 is the value of 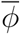 that maximises the solid flux. This expression provides a local carrying capacity for solid transport, with smaller radii and higher *Da* supporting a lower maximum. This flux-limiting behaviour is qualitatively consistent with more detailed suspension models [49].

**FIG. 2.**
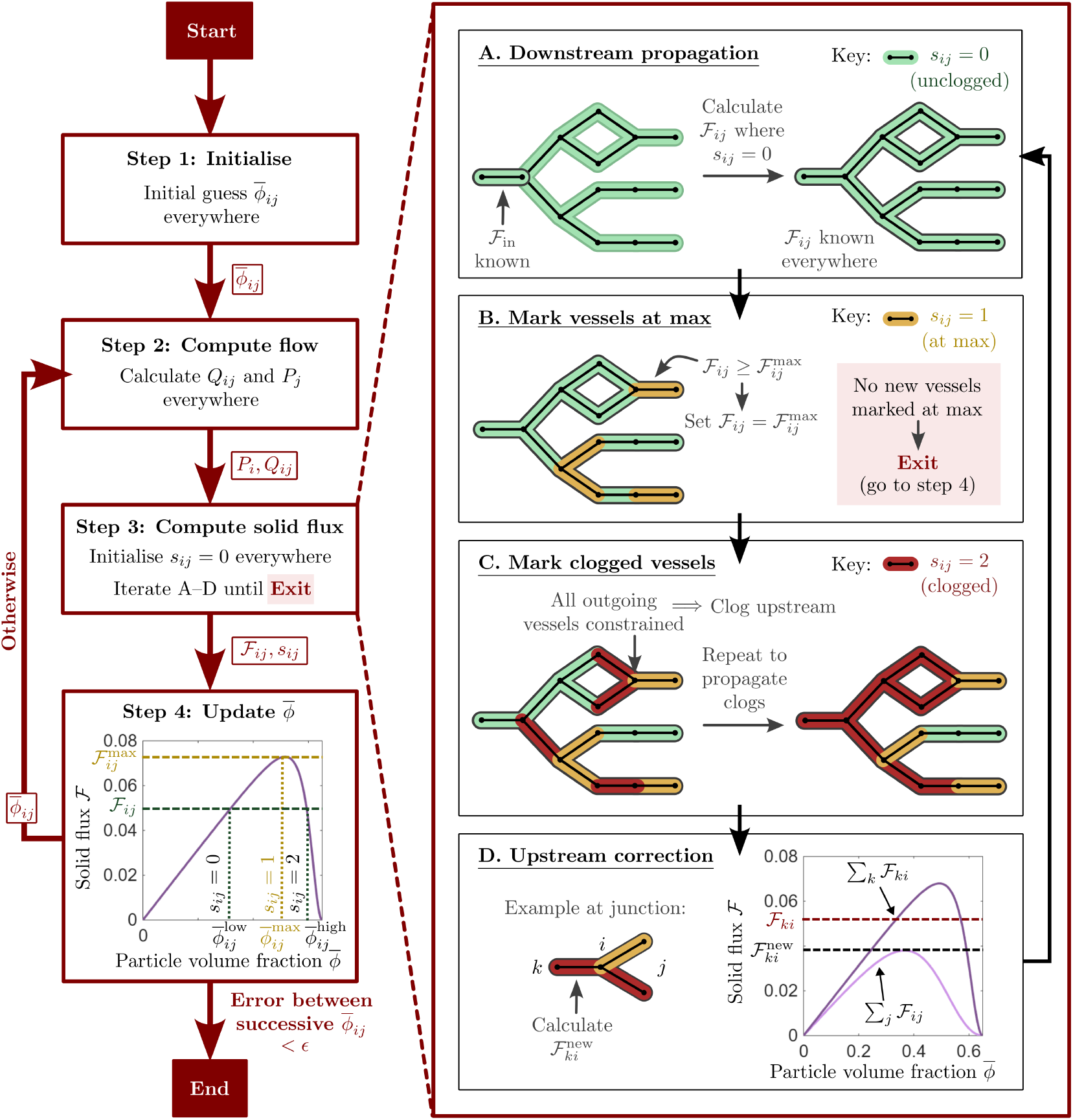
Schematic representation of Algorithm 1. Starting from an initial guess for the particle volume fraction 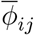, the algorithm iteratively computes the network flow (*Q*_*ij*_), solid flux (ℱ_*ij*_), and 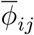 distribution using the vessel constitutive laws (Eq. (1)), mass conservation conditions (Eq. (5)), and solid flux splitting relation (Eq. (6)). Vessel capacity constraints (Eq. (4)) are enforced at each step. When downstream flux limits are exceeded at a junction, the corresponding upstream vessel(s) are marked as clogged and an upstream correction is applied to the solid flux distribution (Eqs. (8-9)).

### C. Flow and transport across junctions: mass conservation, flux splitting, and enforcement of capacity limits

In this work, for simplicity, we restrict our attention to junctions of degree 2 or 3, although similar ideas can be applied to networks with nodes of higher degrees. At each node, we enforce conservation of total flow rate and solid flux. Let *k* and *j* denote the indices of upstream and downstream nodes connected to node *i*, respectively. Then

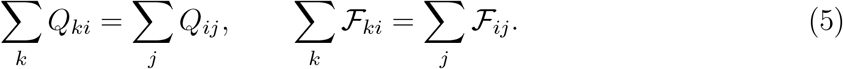

To close the system, we assume that particles are well mixed, and therefore the solid fraction splits in the same way as the total flow via the splitting law

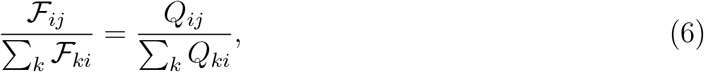

for each *j*. More generally, the framework could accommodate alternative flux partitioning laws accounting for particle deformability or phase separation at bifurcations, as observed for deformable red blood cells in the microvasculature. Such effects are commonly incorporated through empirical or mechanistic hematocrit partitioning laws in blood flow models [52, 53].

To enforce downstream capacity limits arising from each vessel’s maximum capacity 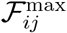 (see Eq. (4)), we set 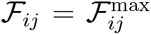 in any outgoing branches where the splitting law causes the flux to exceed its capacity, and, when other outgoing vessels have not reached their flux capacity, redistribute any excess flux among these remaining outgoing branches according to mass balance (Eq. (5)).

### D. Representation of clogging

A capacity violation occurs at a node when the incoming solid flux exceeds the total carrying capacity of all downstream vessels

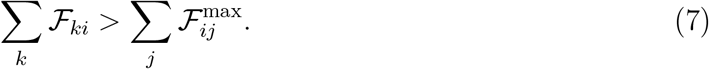

In this case, no distribution of downstream fluxes can simultaneously satisfy mass conservation and the maximum-flux constraints, and so upstream vessels become ‘clogged’ (i.e. they transition to a high-resistance, low-flux state; see Sec. II A). In the model, this ‘clogging’ is represented through an upstream correction to the solid flux distribution and the assignment of clogged vessels to the high particle volume fraction branch of the ℱ_*ij*_ constitutive relation. In this way, capacity violations represent a change in causality, through a change in the direction of information transfer, with the build-up of solid propagating upstream through the network, potentially triggering clogging as it goes.

#### 1. Upstream flux correction

At a junction in which a capacity violation has occurred, the upstream solid flux(es) must be reduced to match the total downstream carrying capacity. For a single upstream vessel at a node *i*, with flux-limited downstream vessel(s), the new solid flux is given via mass balance as

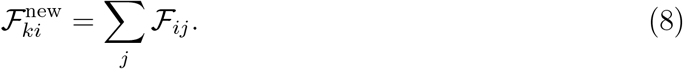

For multiple upstream vessels, the corrected flux is distributed for each *k* via

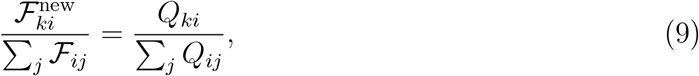

corresponding to a backward application of the splitting law (Eq. 6). Capacity limits are enforced as before, and mass balance is used to redistribute solid flux if this assignment causes either upstream vessel to reach its maximum flux. Physically, this correction reflects the fact that particle accumulation upstream of the junction reduces the rate at which solid material can be transported. A graphical interpretation is shown in Fig. 2.

#### 2. Calculation of 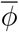 from solid flux

Given the solid flux ℱ_*ij*_ in each vessel, the corresponding particle volume fraction is obtained by inverting the constitutive relation 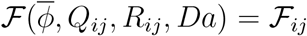. In a vessel *V*_*ij*_, for a given (*Q*_*ij*_, *R*_*ij*_, *Da*) and solid flux 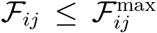, the inversion admits either one or two solutions. When 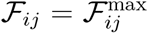, the solution is uniquely given by 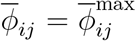. Otherwise, two solutions exist, 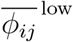 and 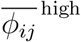, corresponding to lower and higher solutions on either side of 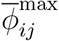 (see Fig. 2).

In unconstrained vessels (vessels which are not clogged or set to their maximum flux), we select the 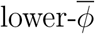 solution. This choice ensures solution continuity: in the absence of flux-limiting constraints, 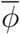 adjusts smoothly to changes in geometry and flow, remaining on the same (lower) branch as the imposed inlet condition. Transition to the upper branch would require a jump in 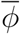, which is not physical in unclogged vessels. This interpretation is consistent with the spatio-temporal solutions of Herale *et al*. [49], which show gradual increases in 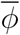 across constrictions when downstream capacity is not exceeded. Throughout this work, the prescribed inlet volume fraction 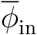 is taken to lie on the lower branch of the flux curve 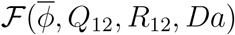 initially.

By contrast, vessels identified as clogged are assigned the upper-branch solution 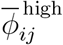, corresponding to a high particle concentration, low-transport state. In this way, particle accumulation arising from downstream capacity constraints is represented by a transition from the lower to upper branch of the constitutive relation.

#### 3. Propagation of clogging

The procedures described above identify clogging following a local capacity violation (Eq.(7)) at a node and represent clogged vessels through the upstream correction and high-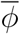 branch selection rules. However, these updates directly affect adjacent upstream junctions, since a vessel marked as clogged at one node serves as a downstream branch for a neighbouring junction. As a result, this can produce junctions for which all downstream vessels are either operating at their maximum solid flux or have become clogged. In such cases, all upstream vessels are also marked as clogged, and the same upstream correction and high-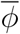 branch selection rules are applied to this newly affected junction. This process is repeated until no additional vessels are marked as clogged, allowing clogging to propagate through the network.

### E. Model initialisation and transport characterisation

In this work, we consider transport in networks with a single inflow vessel and prescribed geometry *{L*_*ij*_, *R*_*ij*_*}* (see Fig. 1e). We set the inlet conditions for the inflow vessel, namely the total inlet flow rate *Q*_in_(= *Q*_12_) and the inlet particle volume fraction 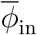. In all simulations, the inlet vessel is scaled such that *Q*_12_, *L*_12_, *R*_12_ = 1. We specify the outlet boundary condition by fixing the pressure *P*_out_ = 0 at all outlet nodes. In this way, the inlet pressure (and total pressure drop Δ*P*_total_) are determined as part of the solution. Along with these boundary conditions, we close the system through prescribing the Darcy number, *Da*, and the maximum packing fraction, *ϕ*_*m*_. This formulation therefore yields a model governed by only a small number of parameters in addition to the prescribed geometry.

To quantify the global impact of clogging on transport, we define the total network resistance as

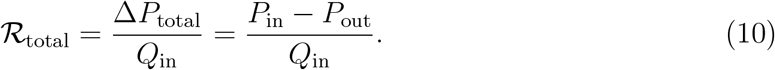

Changes in ℛ_total_ provide a network-level measure of how local flux constraints and clogging alter the ability of the network to transport material.

In addition to the total network resistance, we analyse transport at the vessel scale using the solid fraction ℱ_*ij*_*/Q*_*ij*_, which quantifies the proportion of the total flow that represents solid transport and is therefore central to describing solid flux redistribution and clogging.

## III. NETWORK ALGORITHM

To solve our model for particle flow and transport in a network with specified topology and vessel geometries, we use Algorithm 1 (represented graphically in Fig. 2). The algorithm combines calculation of flow rates and pressures, junction splitting, flux-capacity enforcement, and vessel state-dependent inversion of the solid flux curves, and uses an iterative approach reminiscent of previous methods for flow and hematocrit transport in vascular networks [53]. To track clogging and flux limitation, each vessel is assigned a discrete state *s*_*ij*_ ∈ {0, 1, 2}, where *s*_*ij*_ = 0 corresponds to an unconstrained vessel, *s*_*ij*_ = 1 to a vessel operating at its maximum flux, and *s*_*ij*_ = 2 to a clogged vessel. These states determine solution branch selection for the particle volume fraction as described above. The iterations of Algorithm 1 should not be interpreted as temporal evolution of the suspension. Rather, they form a numerical procedure for finding a steady network configuration that satisfies the constitutive relations, conservation laws, and flux-capacity constraints simultaneously.

### Algorithm 1

Network flow solver with flux-capacity enforcement

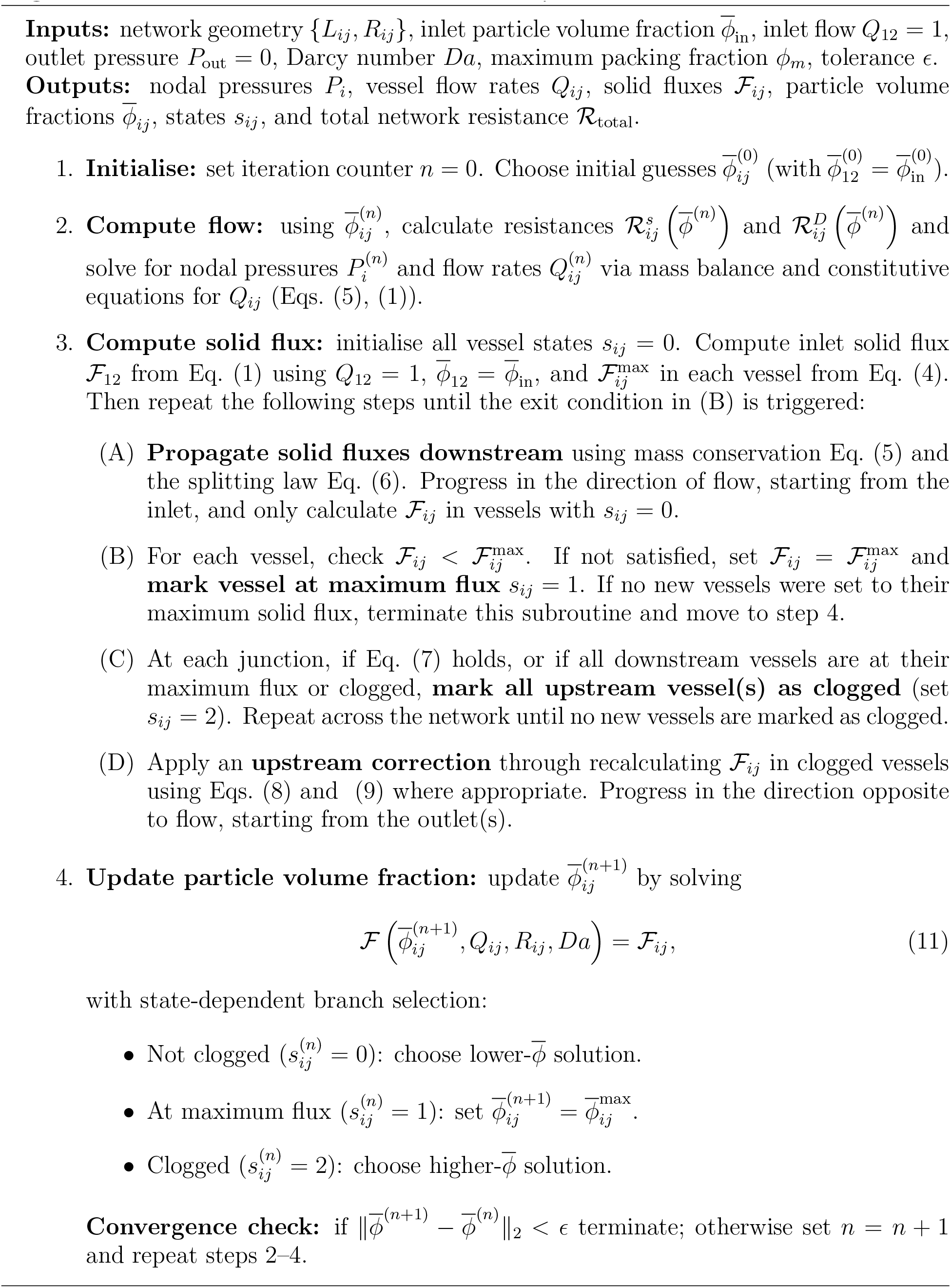

A key component of the method is the nested solid-flux subroutine (see Algorithm 1, Step 3), which enforces vessel flux capacities across the network. Starting from the inlet, fluxes are propagated downstream according to the splitting law (step A), after which local capacity constraints are imposed (step B). When violations occur, vessels are marked at their maximum flux and, if downstream capacity is exceeded, upstream vessels are identified as clogged (step C). Mass conservation is then restored via an upstream correction (step D). This alternating downstream-upstream procedure reflects the directionality of information transport: flux is imposed downstream, while clogging constraints propagate upstream via mass balance. An example of the resulting evolution of vessel states in a representative network is illustrated in Supplementary Video S1. Since no vessel in the network can ever revert from a clogged/maximum flux state to the unclogged state, this subroutine will always terminate after a finite number of iterations.

The non-linear, flux-limited constitutive behaviour introduces hysteresis into the network solution. Because each vessel admits both lower- and higher-branch solutions, multiple steady states may exist for identical input parameters, and the converged solution depends on the initial condition. The choice of initial guess 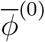 is therefore important for convergence and physical interpretation. Two natural choices are a spatially uniform state 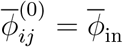, corresponding to a network with all vessels on the lower branch, or a previously converged particle volume fraction distribution, obtained for nearby parameters, which acts as a continuation in parameter space. The latter is particularly appropriate when modelling gradual geometric or parametric changes, as it preserves the history dependence inherent to the model. For the simulations presented here where we will reduce the radius of a vessel to trigger clogging, we combine these approaches. We first compute the steady solution for the network with no reduction in radius using a uniform initial condition 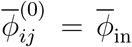. This converged state is then used to initialise further simulations with reduced radii. This biases the solver toward states in which excess solid flux is initially transmitted according to the original geometry, so that any violation of downstream flux limits arises from the imposed geometric changes. This procedure allows the onset and propagation of clogging to be interpreted as a consequence of reductions in radius, rather than as an artefact of initialisation.

For a given initial condition, the solution is obtained by iterating Algorithm 1 until convergence of the particle volume fraction field. Specifically, the outer iteration updates the flow rates, solid fluxes, and particle volume fractions until 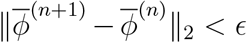. Throughout this work, we use a tolerance of *ϵ* = 10^−6^, for which the algorithm typically converges within 20 iterations. During Step 4, the flux curve 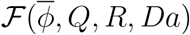 is evaluated numerically on a fine uniform grid (10^5^ values of 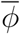), and Eq. (11) is solved by identifying the value of 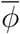 that minimises the error with ℱ_*ij*_, subject to the branch selection criteria.

## IV. RESULTS

To summarise briefly, our non-dimensional model (Sec. II) and numerical algorithm (Sec. III) allow us to simulate the flow rate and particle flux of a particle suspension in a network with prescribed topology and geometry; particle flux limitation in individual vessels is captured by separating the motion of the particle suspension into a particle-rich ‘wet solid’ phase and a Darcy seepage flow of fluid past the particles. Vessel geometry is defined by the radius *R*_*ij*_ and length *L*_*ij*_ of each vessel linking node *i* to node *j*; these are scaled by the dimensional radius and length of the inlet vessel, 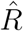 and 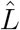, and all vessels in the idealised network motifs considered below have unit dimensionless length (*L*_*ij*_ = 1). As input, we impose the inflow volume fraction 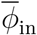, the maximum packing fraction 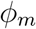, at which particle flux reduces to zero, and the Darcy number 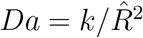, which quantifies the influence of permeability *k* on suspension transport. Our algorithm outputs the flow rate *Q*_*ij*_ and solid flux ℱ_*ij*_ in each vessel (both scaled by the dimensional inflow rate 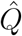), among other quantities (see Algorithm 1). In general, we aim to produce results for a wide range of parameters to reveal the range of possible behaviours in minimal networks. In the main text, the maximum packing fraction is taken to be *ϕ*_*m*_ = 0.85, as a representative value for deformable particles, whose maximum packing fraction can substantially exceed that of rigid spheres [54], for which *ϕ*_*m*_ ≈ 0.65 [55] (see Section IV D for further discussion). For comparison, we repeat the corresponding simulations using *ϕ*_*m*_ = 0.65, with slight adjustments to the network radii to obtain comparable transport behaviour, in Supplementary Figs. S3–S6, giving qualitatively similar results.

### A. Local flux redistribution

We now apply the model to explore the flow of dense particle suspensions in minimal and physiological network topologies. First, we examine how variation in downstream vessel geometry in a three-vessel junction redistributes solid flux between branches, as changes in vessel capacity alter the partitioning of incoming solid flux between downstream vessels, potentially driving one or both branches to reach their maximum solid flux and triggering clogging (which in our framework corresponds to a high-resistance, low-flux state; see Sec. II A). The system consists of one upstream vessel and two downstream vessels, with one downstream radius (*R*_24_) varied while the others are fixed (see schematic in Fig. 3a). For comparison, we also compute a Newtonian reference solution (see Appendix B for details), in which flux capacity constraints and particle accumulation are neglected. This provides a baseline against which the effect of flux limitation on the total network resistance can be examined. Across multiple values of the Darcy number *Da*, we identify three distinct transport states in the 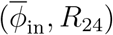 parameter space: an unconstrained state in which all vessels operate below their maximal particle flux 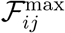, a partially constrained state in which only the vessel with varied radius operates at its maximum, and a fully constrained state in which all vessels are either operating at 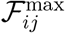 or clogged (Fig. 3a). As *Da* increases (e.g. because of higher permeability), the flux-constrained regions occupy a larger portion of parameter space, because higher *Da* reduces the Darcy resistance 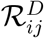 (see Eq. 2) and therefore 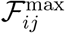, lowering the threshold for flux limitation.

**FIG. 3.**
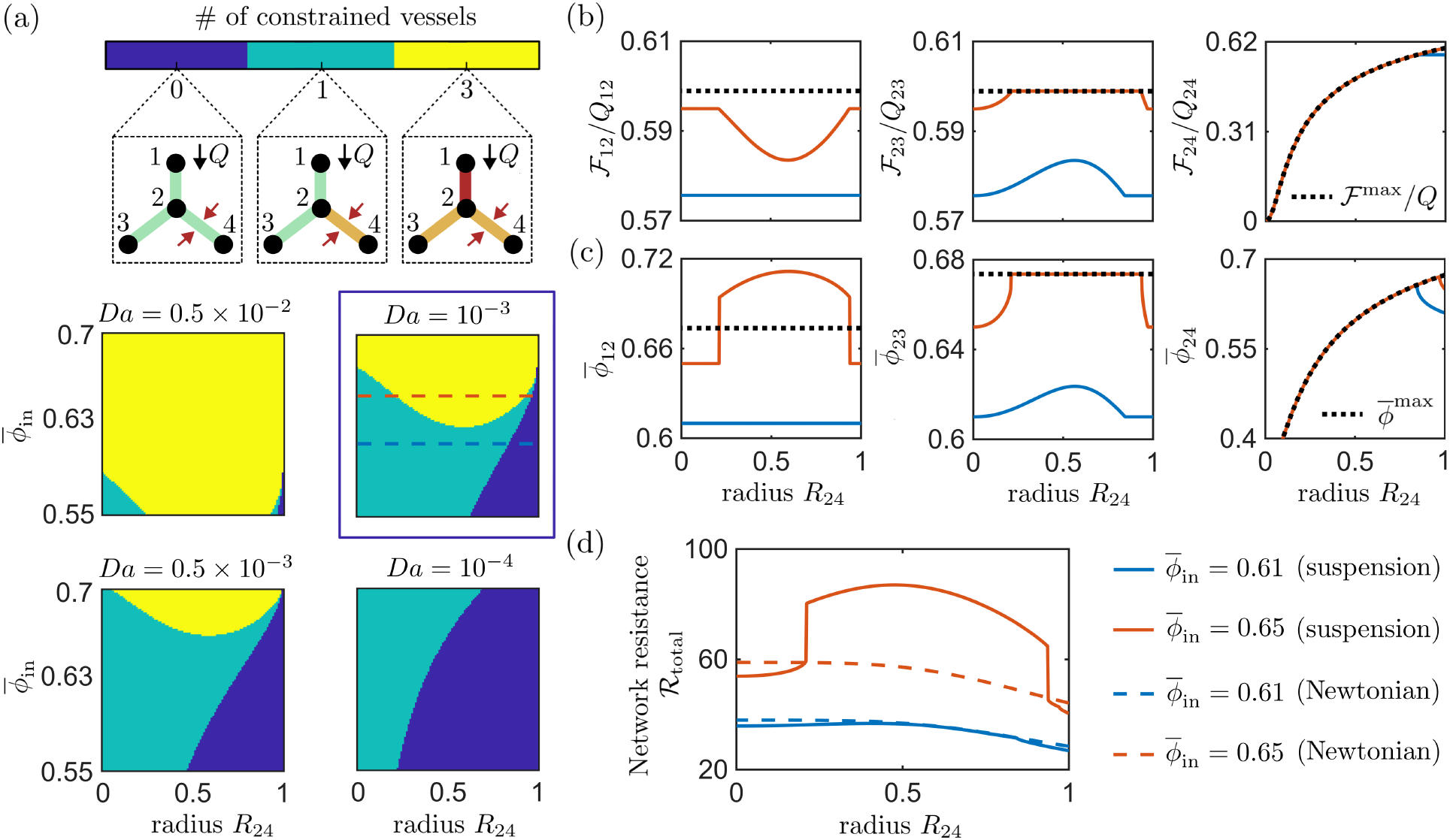
Local redistribution of solid flux in a three-vessel system (as illustrated in panel (a)) as *R*_24_ is varied while the remaining radii are held fixed at *R*_12_ = 1, *R*_23_ = 1. (a) Number of flux-constrained vessels as a function of *R*_24_ and 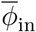, shown for four values of *Da*. Insets illustrate junction states, with unconstrained vessels in green, vessels at their maximum flux in orange and clogged vessels in red. For *Da* = 10^−3^, panels (b–d) show results for two values of 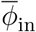 (indicated by dashed lines in (a)): a lower value (blue), for which local redistribution does not produce complete network clogging, and a higher value (red), for which redistribution drives the junction to a fully clogged state. (b) Solid fraction ℱ*/Q* and ℱ^max^*/Q* in each vessel versus *R*_24_ (plots of ℱ and *Q* individually provided in Supplementary Fig. S1). (c) Particle volume fraction 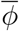 and 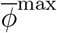 in each vessel versus *R*_24_. (d) Total network resistance ℜ_total_ versus *R*_24_, with the corresponding Newtonian solutions shown with dashed lines. Simulations are shown for *ϕ*_*m*_ = 0.85.

In this minimal network topology, local redistribution of solid flux provides the mechanism by which clogging of the entire network emerges. In the unconstrained regime, the splitting law prescribes constant partitioning of solid flux between the two downstream vessels, and the solid fractions (ℱ_23_*/Q*_23_ and ℱ_24_*/Q*_24_) in both vessels remain constant (Fig. 3b). Correspondingly, the particle volume fraction in the vessel whose radius is varied increases as *R*_24_ decreases, while that in the adjacent downstream vessel remains constant, with both vessels remaining on the lower solution branch of the constitutive relation (Fig. 3c). The network resistance increases gradually as the downstream radius decreases, while remaining comparable to the corresponding Newtonian prediction (Fig. 3d). The onset of local redistribution occurs when the downstream vessel where the radius is varied reaches its maximum solid flux. Once this limit is reached, its solid flux becomes constrained by ℱ^max^, and any excess solid flux must be redirected into the adjacent downstream vessel. Since the maximum solid flux decreases with the vessel radius, an increasing fraction of the incoming solid flux is redirected into the neighbouring vessel. As a result, its solid fraction increases monotonically while its particle volume fraction increases along the lower solution branch, reflecting particle accumulation (Fig. 3b–c). This increase reflects both a redistribution of solid flux into the neighbouring vessel and changes in the underlying flow distribution. The corresponding variations in ℱ and *Q* are shown separately in Supplementary Fig. S1. This regime therefore illustrates the central mechanism of local redistribution: a capacity constraint in one branch alters transport throughout the junction. For sufficiently high 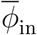, redistribution through continued reduction of the downstream radius eventually drives the adjacent downstream vessel to also reach its maximum solid flux. At this point, neither downstream vessel can accommodate additional solid flux, causing the upstream vessel to become clogged. In this regime, the particle volume fraction in the downstream vessels remains fixed at their maximum values, 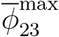 and 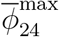, while the inlet vessel transitions from the lower 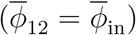 to the upper solution branch (Fig. 3c). The inlet solid fraction ℱ_12_*/Q*_12_ decreases in response to the reduced downstream carrying capacity, and the network resistance exhibits a sharp jump, in contrast to the Newtonian solution which remains smooth (Fig. 3d).

A notable feature of the regime diagram for this minimal network is that further reducing the radius of the downstream vessel can move the system from the fully constrained state to a state in which only that vessel is flux constrained (Fig. 3a). This is because, in the limit of a vanishingly narrow downstream vessel, that vessel’s contribution to solid and interstitial transport becomes negligible, and the junction effectively reduces to a two-vessel system. The redistribution mechanism responsible for overloading the adjacent vessel is therefore removed: all flow is diverted into this neighbouring branch, increasing its total flow (*Q*_23_) relative to solid flux (ℱ_23_) and thereby reducing the solid fraction. This causes this neighbouring vessel to fall below its maximum value and both it and the upstream inlet return to their lower solution branches (Fig. 3b–c). The increase in the inlet solid fraction indicates partial recovery of solid transport through the upstream vessel even while all downstream vessels remain flux constrained, as the reduced influence of the very narrow downstream vessel lowers the severity of the downstream bottleneck. Viewed another way, this demonstrates that the addition of an extra branch to a network can, in fact, promote clogging by enabling redistribution that overloads neighbouring vessels.

### B. Non-local clogging

Having established the role of local flux redistribution in smaller networks, we now demonstrate how non-local clogging emerges in larger branching networks through network-wide redistribution of solid flux (Fig. 4). To isolate this mechanism, we consider a minimal seven-vessel tree where the inlet vessel splits into left- and right-hand branches, each of which further splits into two at the next level (Fig. 4a). The radius *R*_48_ of a downstream vessel located in the lower right-hand branch is varied to impose a downstream flux constraint. We define non-local clogging as the emergence of a clogged vessel not connected to the vessel in which the radius is varied through a continuous chain of flux-constrained vessels (i.e. vessels at maximum flux or clogged). The geometry is chosen such that a vessel in the lower left-hand branch is flux constrained for all parameter values to promote clogging, remaining fixed at its maximum solid flux. All remaining vessels are initially unconstrained (see Fig. 4b).

**FIG. 4.**
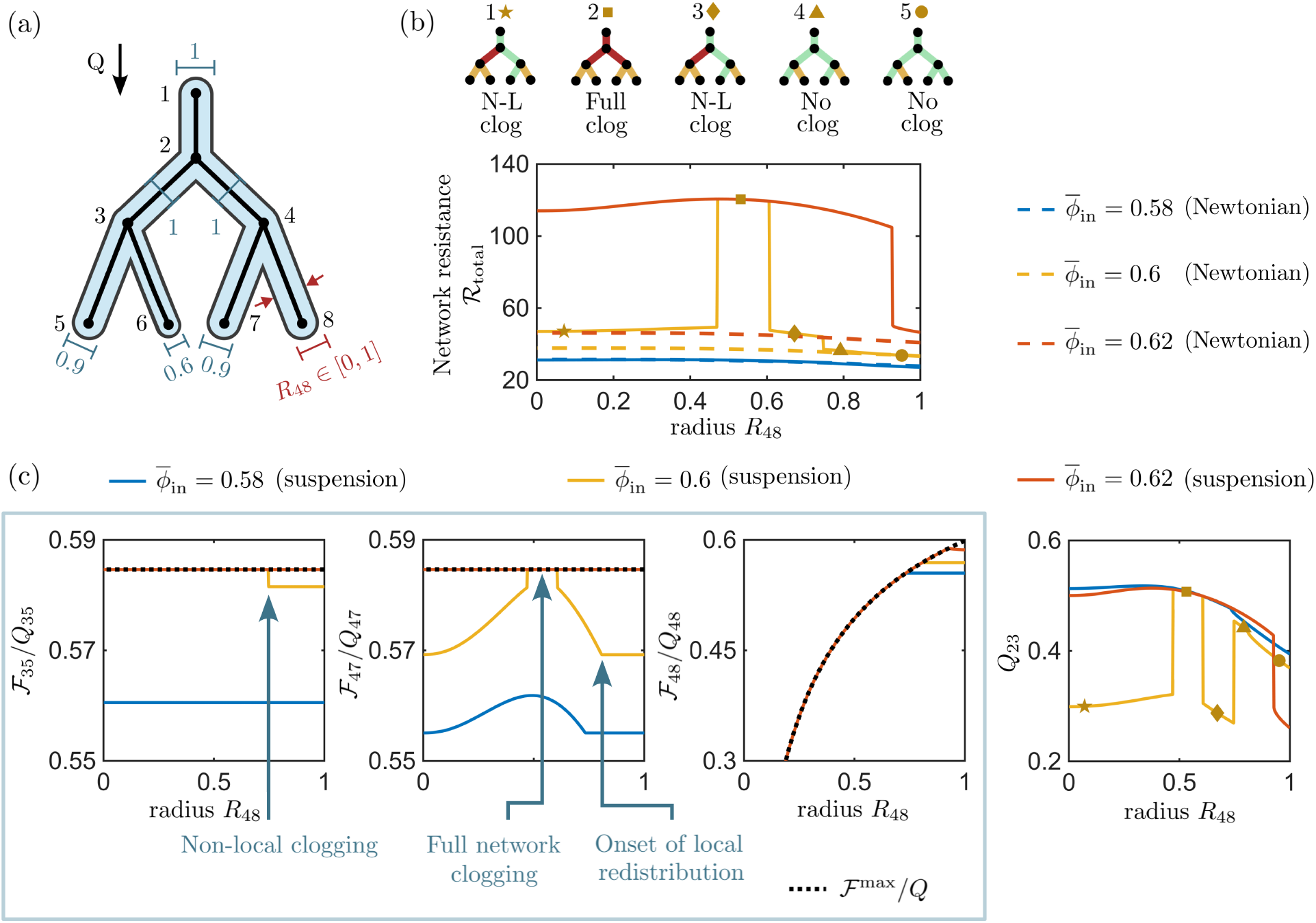
Non-local (N-L) clogging induced through variation in vessel radius in a simple tree network. Non-local clogging is defined as the emergence of a clogged vessel that is not connected to the vessel in which the radius in varied (*V*_48_) through a continuous chain of flux-constrained vessels. The network response is shown for three values of 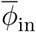: a low value for which neither non-local or full network clogging occurs (blue), an intermediate value for which non-local clogging is followed by full network clogging (yellow), and a high value where the system transitions directly to full network clogging (red). Simulations are performed at *Da* = 10^−3^, *ϕ*_*m*_ = 0.85. (a) Network schematic showing radii. (b) Total network resistance ℛ_total_ over *R*_48_, with the corresponding Newtonian solutions shown with dashed lines. Network schematics 1–5 (left-right) indicate vessel states (unclogged in green, set to maximum flux in orange, clogged in red) for selected values of *R*_48_ for the intermediate 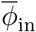 case 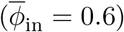. (c) Solid fraction ℱ*/Q* and ℱ^max^*/Q* in three vessels (*V*_35_, *V*_47_, *V*_48_) versus *R*_48_ (left), together with the total flow rate in vessel *V*_23_ (right), illustrating the redistribution of flow associated with non-local clogging.

As the radius *R*_48_ of the downstream vessel is reduced, local redistribution occurs within the right-hand side of the network. When the vessel with reduced radius reaches its maximum solid flux, it becomes flux constrained and excess solid flux is redirected into the adjacent vessel at the same level. Since the maximum solid flux in the constrained lower right-hand vessel decreases as its radius is decreased, an increasing fraction of the incoming solid flux must be accommodated by the adjacent vessel, leading to a monotonic increase in its relative solid flux ℱ_47_*/Q*_47_ (Fig. 4c). This behaviour is analogous to the local redistribution mechanism identified in Sec. IV A.

Non-local clogging emerges when global redistribution of solid flux drives a distant branch to its maximum solid flux, causing clogging upstream. As the downstream capacity on the right-hand side is reduced through further reductions in *R*_48_, the resistance of the right-hand part of the network increases; although the splitting law at the first junction dictates that the solid fractions ℱ*/Q* in the upper left- and right-hand branches remain constant, solid flux is preferentially diverted towards the left-hand side of the network. In contrast to the Newtonian reference solution, this redistribution can overload vessels far from the vessel in which the radius is varied and drive them to their transport limits. For certain values of 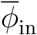, this eventually forces the unconstrained lower left-hand vessel to reach its maximum solid flux, since the adjacent vessel is already flux limited. In this regime, neither vessel on the lower left-hand branch can accommodate additional solid flux, and the upstream vessel transitions to the clogged state (Fig. 4b). This clogging event is non-local: it is triggered by the reduction in radius in the lower right-hand branch rather than a continuous series of upstream clogs. Although the corresponding change in the solid flux fraction in the lower left-hand branch (ℱ_35_*/Q*_35_) appears small in Fig. 4c, this results from simultaneous variations in ℱ_35_ and *Q*_35_, which are larger when considered individually (see Supplementary Fig. S2). The onset of non-local clogging depends on 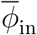: for lower values, the network never reaches the non-local or fully clogged states, whereas for higher values it bypasses the intermediate non-locally clogged state and transitions directly to a fully clogged configuration.

Non-local clogging also does not necessarily persist as the downstream radius *R*_48_ is reduced even further. Following the non-local clogging transition, further reduction in the downstream capacity eventually causes the remaining lower branch on the right-hand side to reach its maximum solid flux, leading to a fully clogged network (Fig. 4b). Then, similarly to the three-vessel case described above, reducing the downstream radius still further can recover the non-locally clogged network state (Fig. 4b). Here the contribution of the narrow lower right-hand vessel to solid and interstitial transport is negligible, and the network behaves as a reduced system in which this vessel is effectively removed, thereby eliminating the redistribution mechanism responsible for the previous transition.

The corresponding changes in flow rate further illustrate the coupling between particle transport and fluid redistribution as the lower right-hand vessel’s radius *R*_48_ is reduced. Prior to non-local clogging, the flow rate in the non-locally clogged branch (*Q*_23_, Fig. 4c) increases as the reduced downstream capacity redirects flow towards this branch. At the onset of non-local clogging, *Q*_23_ decreases abruptly as the newly imposed flux constraints modify the coupled flow and particle distribution, directing more of the suspension flow through the right-hand branch. A further transition to full network clogging results in an abrupt increase in *Q*_23_, which can be interpreted by the fact that the imposed total flux through the network is shared out more equally between the left and right sides of the network in this fully clogged state; the associated network resistance, of course, increases substantially in the clogged state (Fig. 4a). Finally, as the radius of the lower right-hand vessel is reduced still further, we recover the non-locally clogged state, in which there is a second decrease in *Q*_23_. These abrupt changes demonstrate the extent to which clogging transitions are accompanied by reorganisations of both solid transport and fluid flow throughout the network.

### C. Effect of network topology

We next examine how increasing network connectivity alters flux redistribution and clogging behaviour. Starting from the network in Fig. 4, we introduce an additional ‘bypass’ vessel (see Fig. 5), in order to create an alternative route for both fluid and solid transport. This modification increases the connectivity of the network and provides an additional degree of freedom for redistributing solid flux. To assess the impact of the bypass vessel, we vary the radius *R*_48_ of the same downstream vessel as in Fig. 4, which controls the overall transport capacity of the network and sets how close the system is to a flux-limited regime. The effect of the additional pathway depends strongly on this downstream capacity (Fig. 5). When the network is already close to a flux-limited state, the introduction of the bypass vessel can reduce the number of constrained vessels by providing an alternative route for solid flux. However, this effect is non-monotonic: depending on the radius of the bypass vessel, the additional pathway can also redirect flux in a way that triggers constraints elsewhere in the network. For larger downstream capacities, the additional pathway more frequently promotes clogging by modifying the global flow distribution and increasing the solid flux delivered to certain branches. As shown by the representative network states in Fig. 5, this can drive vessels towards their maximum transport capacities, trigger additional flux constraints, and induce upstream clogging that is not present in the original network.

**FIG. 5.**
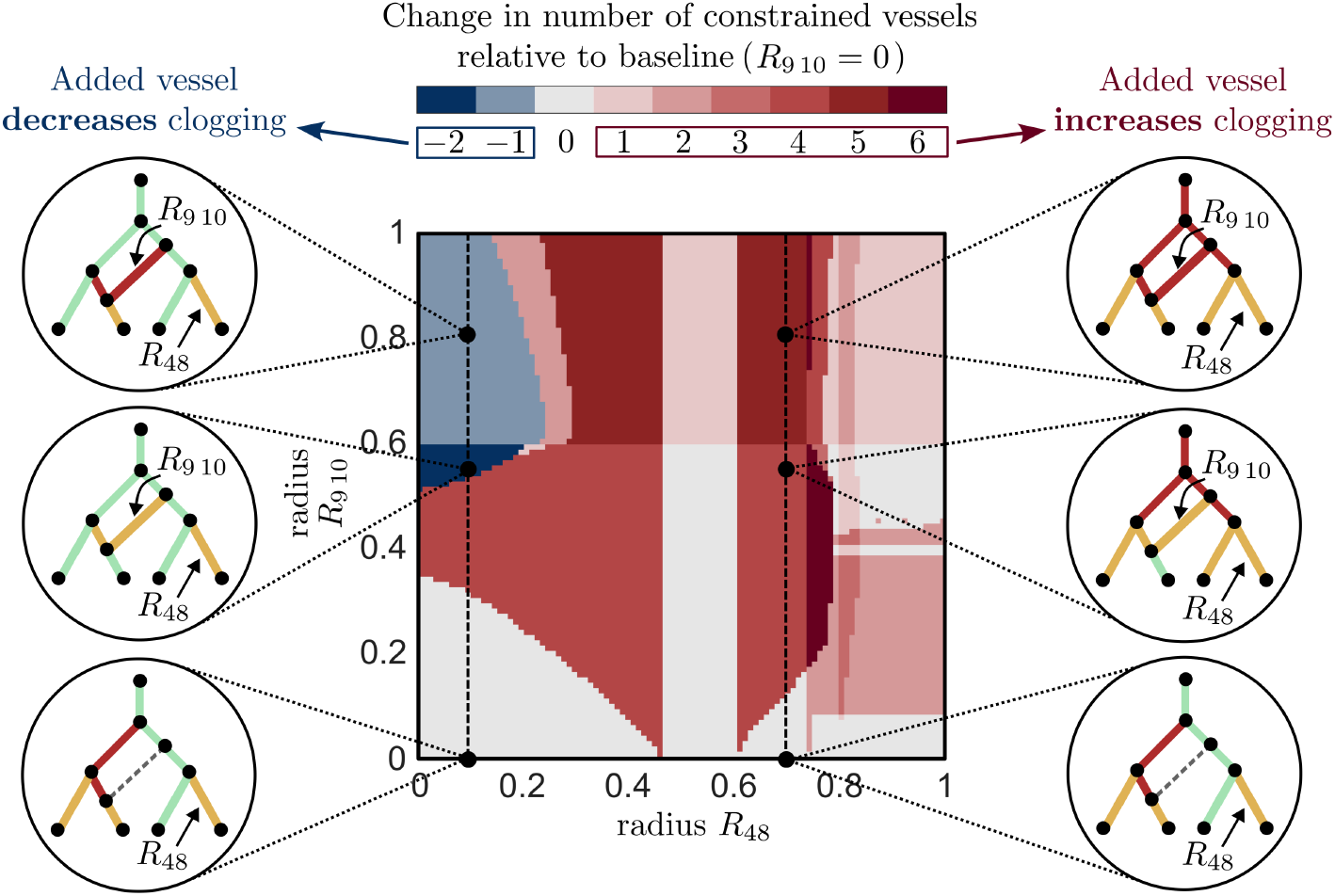
Changing network topology through an additional bypass vessel enhances or reduces clogging depending on downstream capacities. Colormap shows the change in the number of constrained vessels as a function of vessel radii *R*_48_ and *R*_9 10_, relative to the baseline case in which the bypass vessel is removed (*R*_9 10_ = 0). Junction schematics indicate vessel states: unconstrained (green), at maximum flux (orange), and clogged (red). Network geometry is identical to Fig. 4, with additional nodes 9, 10 introduced to split *V*_24_ and *V*_36_ respectively. Results are shown for 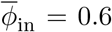, *Da* = 10^−3^, *ϕ*_*m*_ = 0.85. These parameter values correspond to the non-local clogging regime identified in Fig. 4.

These results demonstrate that the impact of increased connectivity depends sensitively on the balance of downstream capacities. Rather than simply relieving bottlenecks, an additional pathway can redistribute fluxes in a way that either reduces or enhances clogging, highlighting the strongly non-local nature of transport in these networks.

### D. Clogging in large-scale physiological networks

Finally, as a proof of concept, we predict how the mechanisms identified above could manifest at the network scale in physiological contexts, by applying the solution algorithm to vascular arterial tree networks derived from retinal vasculature geometries [31] (Fig. 6). In contrast with the idealised motifs considered above, vessel radii and lengths are taken directly from the generated network geometry. The resulting network contains a single inlet. As a simple representation of pathological occlusion in a retinal network, we explore the implications of reducing a short vessel segment – chosen arbitrarily but with approximate similarities to the vessels occluded to represent diabetic retinopathy in [31] – to 1% of its original radius, thereby substantially decreasing its maximum sustainable solid flux.

**FIG. 6.**
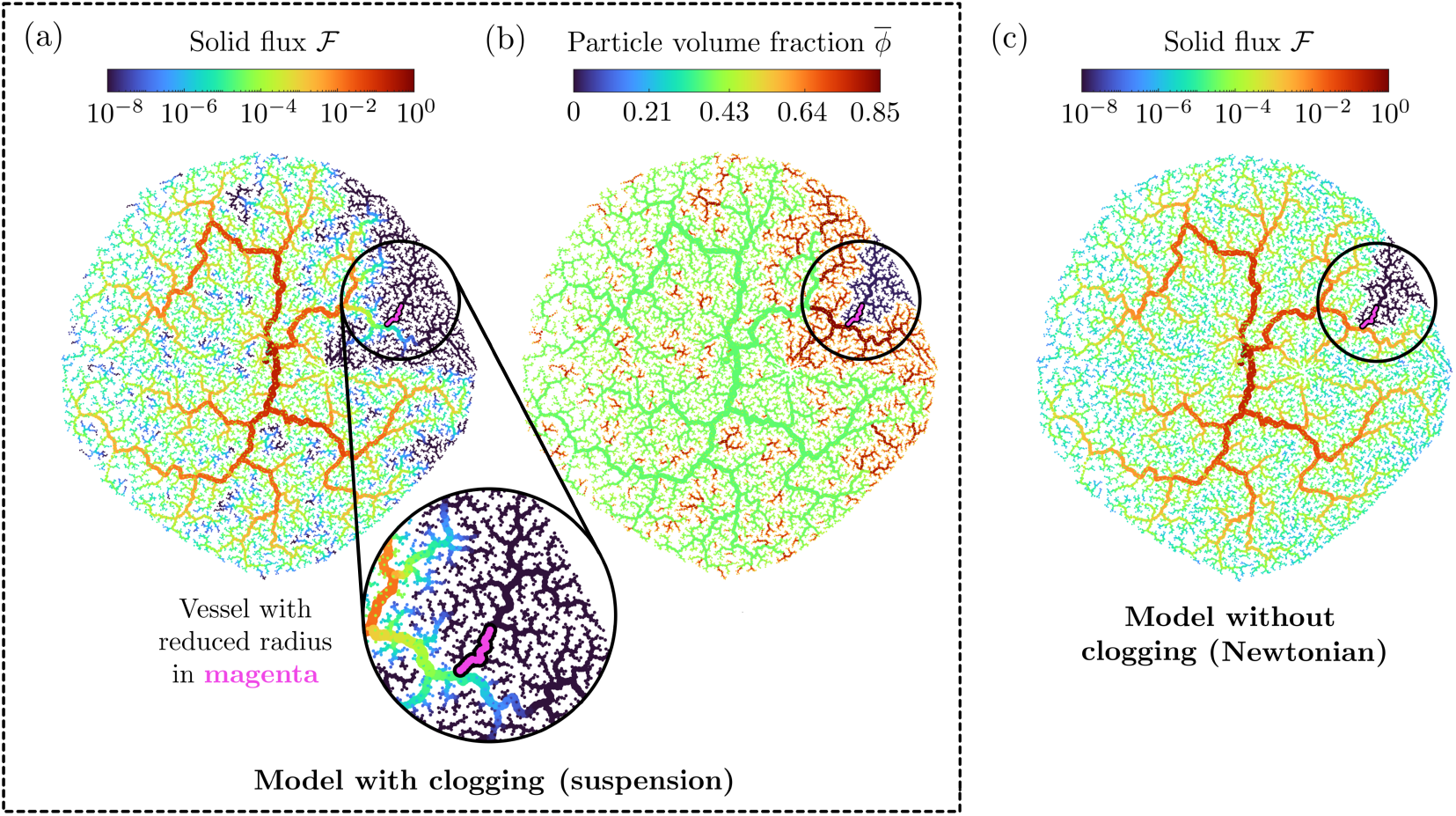
Transport in a complex network (geometry data from [31]) with a short vessel segment (magenta) reduced to 1% of its original radius. (a) Solid flux distribution ℱ and (b) particle volume fraction 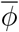 for the suspension model where 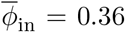, *ϕ*_*m*_ = 0.85 and *Da* = 10^−5^. (c) Proxy for solid flux in a Newtonian fluid, defined as 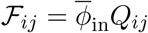, plotted for 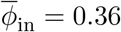.

We perform minimal simulations of retinal blood flow as follows. We set the inlet particle volume fraction to 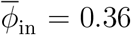, corresponding to the lower end of the physiological range of adult haematocrit (0.36–0.5) [56]. For dense suspensions of red blood cells (RBCs), estimates of the maximum packing fraction depend on factors such as cell shape and deformability, and have been estimated to range from *ϕ*_*m*_ ≈ 0.6 for artificially hardened RBCs to *ϕ*_*m*_ ≈ 1 for healthy, deformable RBCs [54]. Here, we therefore adopt the intermediate value *ϕ*_*m*_ = 0.85 as a representative maximum packing fraction for dense red blood cell suspensions with a pathologically reduced deformability. For simplicity, we do not account for other more complex effects of RBC deformability, such as plasma skimming [52, 53], or for localised reductions in RBC deformability that could occur in oxygen-depleted regions of the retina in sickle cell disease. To estimate the Darcy number, we adopt the commonly used Kozeny-Carman scaling 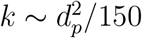, where *d*_*p*_ ≈ 8 *µ*m is the RBC diameter [57], giving *k* ~ 10^−13^ m^2^. The inlet vessel corresponds to the central retinal artery, where reported radii lie in the range 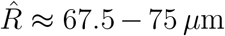 [31]. This yields a Darcy number 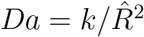 of order 10^−5^, and we therefore adopt *Da* = 10^−5^ in these simulations.

To highlight the effect of flux-capacity limitations, we again compare the suspension model with a Newtonian reference solution in which solid transport is defined by 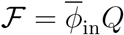 and is unconstrained by flux-capacity limitations (see Appendix B). The suspension and Newtonian models respond very differently to the segment with reduced vessel radius. Both models exhibit reduced downstream transport in response to the reduced vessel radius. In the Newtonian reference case, the upstream flux distribution follows the expected Newtonian scaling, with flow partitioned smoothly at junctions according to vessel radii/resistances (Fig. 6c). By contrast, the suspension model exhibits significant spatial heterogeneity in solid flux with regions throughout the network in which flux is reduced (Fig. 6a). These regions correspond to clogged vessels in our framework, associated with high 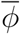 (Fig. 6b). Consistent with the behaviour predicted from the simple motifs in Sec. IV B, the resulting clogging dynamics are strongly non-local, and influence transport far from the vessel segment in which the radius has been reduced. Simulations with different values of *ϕ*_*m*_ show that the maximum packing fraction also controls the degree of clogging observed (Supplementary Fig. S6): a lower *ϕ*_*m*_, corresponding approximately to hardened RBCs, leads to widespread clogging, whereas setting *ϕ*_*m*_ = 1, corresponding approximately to healthy deformable RBCs, suppresses network-level clogging.

These results demonstrate that the local and non-local mechanisms identified in simple motifs extend to complex networks, where they collectively generate large-scale heterogeneity in solid transport. Notably, the emergence of widespread transport suppression does not require explicitly time-dependent dynamics or more sophisticated rheological constitutive laws. Rather, the interplay between finite flux constraints, flow redistribution, and network connectivity alone is sufficient to produce complex network-scale transport patterns. Although these network geometries are relevant to biological transport networks, we emphasise that these proof-of-concept results illustrate physical mechanisms but do not provide a detailed physiological description of a specific disease at this stage.

## V. DISCUSSION

We have built a minimal mechanistic model that combines dense-suspension rheology with network-scale flux redistribution rules to predict clogging over networks. Our model exhibits vessel-level solid flux limits, which are motivated through flux curves that depend on vessel radius and Darcy number, which captures the ease of interstitial fluid flow past particles in the suspension. Through enforcing these limits via junction-level rules which are embedded within a global fluid solver, the model captures how local reductions in radius generate upstream accumulation and sharp resistance increases through a transition in particle volume fraction 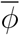 from a lower to an upper solution branch. Analysis of simple motifs reveals that clogging can also be triggered in adjacent vessels, non-locally across the network, through redistribution of solid flux, providing a potential mechanistic explanation for experimentally observed cascading and cooperative blockage in microfluidic and porous networks [58, 59].

We have also demonstrated that the addition of pathways that increase network connectivity does not necessarily improve transport in flux-limited networks. Depending on existing downstream constraints, a new vessel can either relieve clogging by redistributing solid flux or promote it by driving previously unconstrained vessels towards their maximum flux. This behaviour is reminiscent of a classic feature of traffic flow known as Braess’ paradox, in which adding an additional route to a road network can reduce overall performance by redistributing traffic and increasing the utilisation of some routes despite the increased connectivity [60]. Such effects are not limited to traffic networks: recent experiments in microfluidic networks have demonstrated a fluid-mechanical analogue of Braess’ paradox, where modifying network connectivity can decrease the total flow rate for a fixed pressure drop due to non-linear flow-pressure relationships in individual channels, achieved using obstacles that induce inertial flow recirculation [61]. Here, we predict that particle suspension flow provides a natural non-linear pressure-flow relationship owing to flux limitation and suspension rheology; the resultant network behaviour exhibits a similar fluid analogue of Braess’ paradox.

Applied to a bio-inspired network, the model predicts spatial heterogeneity in solid flux and non-local clogging in certain parameter regimes, in contrast to Newtonian flow models. Broadly, this offers a physical route by which small-scale obstructions could lead to large-scale dysfunction in biological transport systems. In vascular systems, for example, a localised occlusion may alter red blood cell transport and promote secondary clogging elsewhere in the network, especially in pathological scenarios with reduced red blood cell deformability. Similar redistribution-driven effects may also arise in plant vasculature, microbial transport networks, and other biological systems in which transport occurs close to local carrying-capacity limits [32, 62, 63]. More widely, this behaviour is also related to redistribution-driven failures in electrical power grids [64], where local failures redistribute power flow and can overload transmission lines with finite carrying capacities, triggering cascading failures. In our model, redistribution of solid flux can similarly overload vessels with finite transport capacities, producing secondary clogging elsewhere in the network.

In our framework, it is convenient to alter the rheological model of suspension flow in individual vessels: the network-wide algorithms would remain unchanged, and only the local constitutive equations for flow rate and particle flux would require updating. This makes it straightforward to incorporate more detailed rheological models [49], and additional physical effects such as particle slip at vessel walls [65, 66], or experimentally measured flux curves and transport laws obtained directly from specific systems. Our results complement particle-resolved and stochastic approaches [44, 45] by providing a computationally efficient, network-level description of transport and clogging. The present model focuses on steady transport in acyclic networks, providing a minimal setting in which the consequences of flux limitation can be studied. Future extensions could explore cyclic network topologies, time-dependent clog growth, and more detailed constitutive descriptions of dense suspension transport.

To summarise, our results demonstrate that finite solid transport capacities can qualitatively alter transport of dense particle suspensions through networks, introducing non-local interactions between branches and making network performance dependent on flux limits in vessels and the redistribution of particles, in addition to vessel resistances. By linking vessel-scale suspension physics to network-scale redistribution and transport failure, the present framework provides a minimal continuum description of how clogging emerges and propagates in complex flow networks. We anticipate that these ideas will be relevant across a wide range of biological, geophysical, and engineered systems in which particle transport occurs near local carrying capacity limits.

## Acknowledgements

We acknowledge helpful discussions with Ranjan Rajendram and Benjamin Walker. We also acknowledge Simon Walker-Samuel for helpful discussions and for providing the physiological network geometry. P.P. was supported by a UK Research and Innovation (UKRI) Future Leaders Fellowship [MR/V022385/1, UKRI2746].

## Data Accessibility

All data underlying the conclusions of the paper is provided in the paper. Code to generate the results in this paper can be found at https://github.com/caravn15/suspension-clogging-networks, where the physiological network geometry is also provided. The network geometry was generated using RetinaGen, which is freely available at https://github.com/simonwalkersamuel/retinasim (see [31] for details).

## Appendix A Appendix: single-vessel suspension model

Using a simplified version of the model proposed by Herale *et al*. [49], we define the constitutive relations applied to each vessel (edge) of the network.

### 1. Conservation of mass and momentum, and constitutive behaviour

Consider a straight cylindrical vessel of radius *R* and length *L* aligned with the *z* direction, with axisymmetric flow driven by a pressure gradient *G >* 0. The suspension consists of solid particles occupying a volume fraction *ϕ* and interstitial fluid occupying a fraction 1 − *ϕ*. The overall axial component of the velocity of the suspension *U* can be written in terms of the solid and fluid axial velocities *u*_*s*_ and *u*_*f*_ via *U* = *ϕu*_*s*_ + (1− *ϕ*)*u*_*f*_. Assuming each phase is individually incompressible, conservation of mass yields

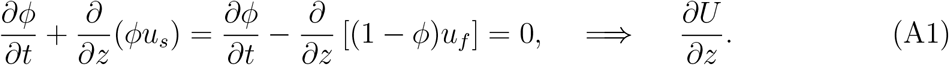

It is more convenient (see [49, 67]) to decompose *U* into the velocity of two phases that track the motion of solid particles and fluid moving at the same speed, and the differential transport of fluid through these moving particles, respectively, via *U* = *u*_*s*_ + *u*_*D*_, where *u*_*s*_ = *ϕu*_*s*_ + (1 − *ϕ*)*u*_*s*_ is the speed of the ‘wet solid’ phase, and *u*_*D*_ ≡ (1 − *ϕ*)(*u*_*f*_ − *u*_*s*_) the speed of the differential ‘Darcy’ flow.

For unidirectional, steady flow, momentum balance for the suspension yields

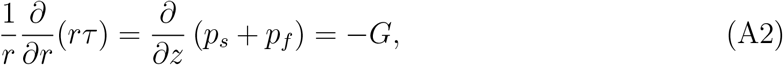

where *τ* = *τ*_*rz*_ is the only non-zero component of the deviatoric stress tensor of the suspension. The total pressure is broken down into the fluid pressure *p*_*f*_ and the excess ‘particle pressure’ *p*_*s*_ that arises due to the interactions of the solid particles. Eq. (A2) is integrated to give a linear shear-stress profile across the pipe, *τ* = −*Gr/*2.

Following [49, 68], the suspension shear stress and particle pressure are related to the local shear rate via *ϕ*-dependent shear and normal viscosities, with *τ* = *η*_*f*_ *η*_*s*_(*ϕ*)∂*u*_*s*_*/*∂*r* and *p*_*s*_ = *η*_*f*_ *η*_*n*_(*ϕ*) |∂*u*_*s*_*/*∂*r*|, where *η*_*f*_ is the fluid viscosity and

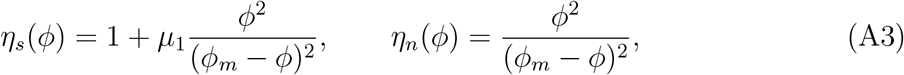

with *µ*_1_ a constant static friction coefficient that we set to unity for the whole of this work. These laws capture the divergence of viscosity as the solid fraction approaches its maximum value *ϕ* → *ϕ*_*m*_; they are a slightly simplified, but qualitatively similar, form to those measured by [68]. It follows immediately that for steady flow in a straight pipe, *p*_*s*_ cannot vary along the pipe and ∂*p*_*s*_*/*∂*z* = 0, meaning ∂*p*_*f*_ */*∂*z* = −*G* in (A2).

The differential fluid flow through the suspension is governed by Darcy’s law,

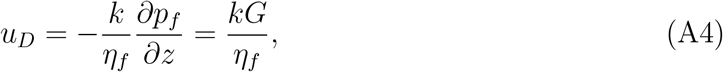

where *k*(*ϕ*) describes the permeability of the suspension.

### 2. Pipe-averaging and flux relations

The steady pipe-flow equations comprise expressions for the total flux *Q* of the suspension and the flux ℱ of the particle alone,

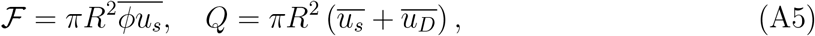

where we use the notation 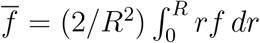 to denote the average across the pipe. The associated pressure drop along the pipe is 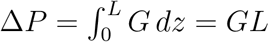.

To proceed, we make the assumption that 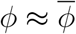, which dramatically simplifies the algebraic expressions compared with those of [49], without changing the qualitative behaviour. Given this assumption, one can treat the suspension as a generalised Newtonian fluid with viscosity 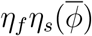, and simply integrate (A2) to find the solid velocity, imposing no slip (*u*_*s*_ = 0) on the pipe walls. The fluxes in (A5) follow from further integration, and can be written as

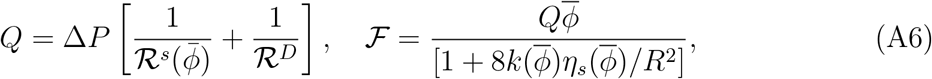

where 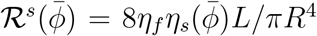 and ℛ^*D*^ = *η*_*f*_ *L/πkR*^2^ are the suspension and Darcy resistances, which effectively act in parallel to make up the total resistance of the suspension in the pipe.

In the expression for ℱ in (A6) we have eliminated Δ*P* in favour of *Q*. From this expression it is clear that 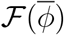 is a non-monotonic function, if the other parameters are held fixed, with ℱ → 0 for both 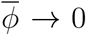 and 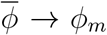, given the form of *η*_*s*_ in (A3). This behaviour implies the existence of a maximum value of ℱ at an intermediate solid fraction, as discussed in the main text.

### 3. Non-dimensionalisation

We scale the system using the properties of the inlet vessel for the network: that is, we scale radial lengths with the inlet radius 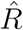, axial lengths with the inlet length 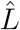, fluxes with the imposed inlet flux 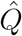, and pressures with 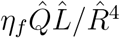. The resulting dimensionless constitutive relations for the fluxes *Q*_*ij*_ and ℱ_*ij*_ in vessel *V*_*ij*_ are

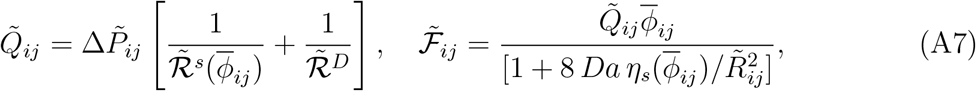

where the dimensionless resistances are 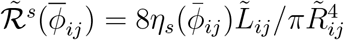 and 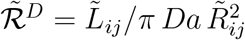, with 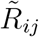 and 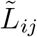 the dimensionless radius and length of vessel *V_ij_*, respectively. The Darcy number 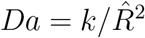 compares the permeability — which scales with the square of the typical particle size — with the square of the characteristic pipe radius 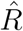. It should therefore be significantly smaller than unity for a continuum description of the suspension to be reasonable. We make the further simplifying assumption that the permeability does not vary appreciably with 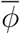, such that *Da* is a constant (again, relaxing this assumption increases algebraic complexity without significantly changing qualitative behaviour).

Throughout the main manuscript, dimensionless quantities are denoted without tildes. These constitutive relations provide the local transport laws from which the network-scale redistribution and clogging behaviour studied in the main text emerge.

## Appendix B Newtonian reference solution

Throughout the results section, to provide a baseline for comparison, we compute a Newtonian reference solution on the same network geometry as the suspension model. This reference solution isolates the effects of network geometry and flow distribution from those arising from suspension rheology and flux limitation.

To enable a direct comparison with the suspension model, the Newtonian fluid is assigned a constant viscosity equal to the suspension viscosity evaluated at the prescribed inlet particle volume fraction, 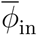. Therefore, each vessel contains a fluid of viscosity 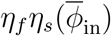, where the functional form of *η*_*s*_ is given in Eq.(A3). The dimensional resistance of vessel *V*_*ij*_ is

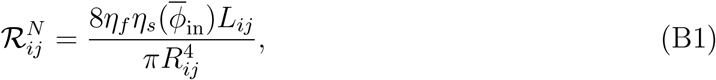

and nodal pressures and flow rates are obtained by solving the network mass conservation equations (Eq. (A1)) together with the constitutive relation

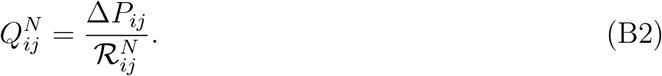

Using the same non-dimensionalisation as in Appendix A, with pressure scaled by 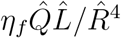, flow rate by 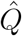, lengths by 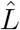, and radii by 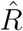, the dimensionless resistance becomes

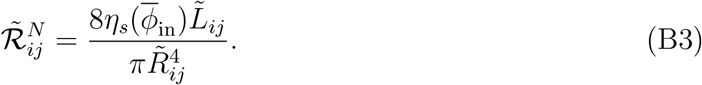

Unlike the suspension model, no flux-capacity constraints, particle accumulation, or branch selection rules are imposed. Particle transport is assumed to behave as a passive tracer with uniform concentration equal to the inlet value 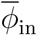. The corresponding dimensionless solid flux in any vessel is therefore simply

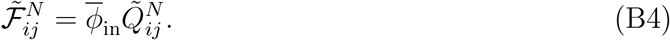

## Notes

https://github.com/caravn15/suspension-clogging-networks

### Competing Interest Statement

The authors have declared no competing interest.

## References

[1] N. Tufenkji and M. Elimelech, Correlation equation for predicting single-collector efficiency in physicochemical filtration in saturated porous media, Environ. Sci. Technol 38, 529 (2004).

[2] J. T. Santini and M. J. Cima, A controlled-release microchip, Nature 397, 335 (1999).

[3] S. Isogai, M. Horiguchi, and B. M. Weinstein, The vascular anatomy of the developing zebrafish: An atlas of embryonic and early larval development, Dev. Biol. 230, 278 (2001).

[4] K. J. Richardson, L. Kuck, and M. J. Simmonds, Beyond oxygen transport: active role of erythrocytes in the regulation of blood flow, Am. J. Physiol. Heart Circ. Physiol. 319, H866 (2020).

[5] B. Choat, R. Munns, M. McCully, J. Passioura, S. Tyerman, H. Bramley, and M. Canny, Water movement in plants, in Plants in Action, edited by B. Choat and R. Munns (Australian Society of Plant Scientists, Melbourne, Australia, 2010) Chap. 3.

[6] K. H. Jensen, K. Berg-Sørensen, H. Bruus, N. M. Holbrook, J. Liesche, A. Schulz, M. A. Zwieniecki, and T. Bohr, Sap flow and sugar transport in plants, Rev. Mod. Phys. 88, 035007 (2016).

[7] W. Konrad and A. Roth-Nebelsick, The phloem: A case of theory against experiment, J. Geophys. Res. Biogeosci. 128, e2023JG007508 (2023).

[8] L. Boddy, J. Hynes, D. P. Bebber, and M. D. Fricker, Saprotrophic cord systems: dispersal mechanisms in space and time, Mycoscience 50, 9 (2009).

[9] A. Tero, S. Takagi, T. Saigusa, K. Ito, D. P. Bebber, M. D. Fricker, K. Yumiki, R. Kobayashi, and T. Nakagaki, Rules for biologically inspired adaptive network design, Science 327, 439 (2010).

[10] L. L. M. Heaton, E. López, P. K. Maini, M. D. Fricker, and N. S. Jones, Advection, diffusion, and delivery over a network, Phys. Rev. E 86, 021905 (2012).

[11] E. C. Hammer, C. Arellano-Caicedo, P. M. Mafla-Endara, E. T. Kiers, T. Shimizu, P. Ohlsson, and K. Aleklett, Hyphal exploration strategies and habitat modification of an arbuscular mycorrhizal fungus in microengineered soil chips, Fungal Ecol. 67, 101302 (2024).

[12] E. Katifori, G. J. Szöllősi, and M. O. Magnasco, Damage and fluctuations induce loops in optimal transport networks, Phys. Rev. Lett. 104, 048704 (2010).

[13] K. Alim, G. Amselem, F. Peaudecerf, M. P. Brenner, and A. Pringle, Random network peristalsis in physarum polycephalum organizes fluid flows across an individual, PNAS 110, 13306 (2013).

[14] E. Katifori, The transport network of a leaf, C. R. Phys. 19, 244 (2018).

[15] H. Ronellenfitsch and E. Katifori, Phenotypes of vascular flow networks, Phys. Rev. Lett. 123, 248101 (2019).

[16] A. F. van Tol, A. Roschger, F. Repp, P. Schneider, K. Klaushofer, and P. Fratzl, Network architecture strongly influences the fluid flow pattern through the lacunocanalicular network in human osteons, Biomech. Model. Mechanobiol. 19, 823 (2020).

[17] N. J. Karst, J. B. Geddes, and R. T. Carr, Model microvascular networks can have many equilibria, Bull. Math. Biol. 79, 662 (2017).

[18] T. Gavrilchenko and E. Katifori, Resilience in hierarchical fluid flow networks, Phys. Rev. E 99, 012321 (2019).

[19] E. Dressaire and A. Sauret, Clogging of microfluidic systems, Soft Matter 13, 37 (2017).

[20] B. Dincau, E. Dressaire, and A. Sauret, Clogging: The self-sabotage of suspensions, Phys. Today 76, 24 (2023).

[21] G. C. Agbangla, P. Bacchin, and E. Climent, Collective dynamics of flowing colloids during pore clogging, Soft Matter 10, 6303 (2014).

[22] A. Marin and M. Souzy, Clogging of noncohesive suspension flows, Annu. Rev. Fluid Mech. 57, 89 (2025).

[23] P. Bacchin, P. Aimar, and R. Field, Critical and sustainable fluxes: Theory, experiments and applications, J. Membr. Sci. 281, 42 (2006).

[24] L. Pang, S. Shen, C. Ma, T. Ma, R. Zhang, C. Tian, L. Zhao, W. Liu, and J. Wang, Deformability and size-based cancer cell separation using an integrated microfluidic device, Analyst 140, 7335 (2015).

[25] H. M. Wyss, D. L. Blair, J. F. Morris, H. A. Stone, and D. A. Weitz, Mechanisms for clogging of microchannels, Phys. Rev. E 74, 061402 (2006).

[26] T. W. Secomb, Blood flow in the microcirculation, Annu. Rev. Fluid Mech. 49, 443 (2017).

[27] A. S. Popel and P. C. Johnson, Microcirculation and hemorheology, Annu. Rev. Fluid Mech. 37, 43 (2005).

[28] G. R. Cokelet and H. J. Meiselman, Rheological properties of human blood and plasma, Handbook of Hemorheology and Hemodynamics, 45 (2007).

[29] F. J. Meigel and K. Alim, Flow rate of transport network controls uniform metabolite supply to tissue, J. R. Soc. Interface 15, 20180075 (2018).

[30] X. Lu, M. M. Galarneau, J. M. Higgins, and D. K. Wood, A microfluidic platform to study the effects of vascular architecture and oxygen gradients on sickle blood flow, Microcirculation 24, e12357 (2017).

[31] E. E. Brown, A. A. Guy, N. A. Holroyd, et al., Physics-informed deep generative learning for quantitative assessment of the retina, Nat. Commun. 15, 6859 (2024).

[32] K. Drescher, Y. Shen, B. L. Bassler, and H. A. Stone, Biofilm streamers cause catastrophic disruption of flow with consequences for environmental and medical systems, PNAS 110, 4345 (2013).

[33] J. Kim, H. Choi, and Y. A. Pachepsky, Biofilm morphology as related to the porous media clogging, Water Res. 44, 1193 (2010).

[34] J. Maruyama, P. R. Juvvadi, K. Ishi, and K. Kitamoto, Three-dimensional image analysis of plugging at the septal pore by woronin body during hypotonic shock inducing hyphal tip bursting in the filamentous fungus aspergillus oryzae, Biochem. Biophys. Res. Commun. 331, 1081 (2005).

[35] E. J. Diamantopoulos, C. Kittas, D. Charitos, M. Grigoriadou, G. Ifanti, and S. A. Raptis, Impaired erythrocyte deformability precedes vascular changes in experimental diabetes mellitus, Horm. Metab. Res. 36, 142 (2004).

[36] S. K. Ballas, Sickle cell disease: Classification of clinical complications and approaches to preventive and therapeutic management, Clin. Hemorheol. Microcirc. 68, 105 (2018).

[37] M. Tsai, A. Kita, J. Leach, R. Rounsevell, J. N. Huang, J. Moake, R. E. Ware, D. A. Fletcher, and W. H. Lam, In vitro modeling of the microvascular occlusion and thrombosis that occur in hematologic diseases using microfluidic technology, J. Clin. Invest. 122, 408 (2012).

[38] S. S. Shevkoplyas, T. Yoshida, S. C. Gifford, and M. W. Bitensky, Direct measurement of the impact of impaired erythrocyte deformability on microvascular network perfusion in a microfluidic device, Lab Chip 6, 914 (2006).

[39] T. Y. Wong and I. U. Scott, Retinal-vein occlusion, N. Engl. J. Med. 363, 2135 (2010).

[40] A. Bhattacharya, Predicting retinal haemorrhage following retinal vein occlusion, Ph.D. thesis, University of Glasgow (2026).

[41] P. A. Roberts, E. A. Gaffney, P. J. Luthert, A. J. E. Foss, and H. M. Byrne, Mathematical and computational models of the retina in health, development and disease, Prog. Retin. Eye Res. 53, 48 (2016).

[42] A. J. Parry and O. Millet, Modeling blockage of particles in conduit constrictions: Dense granular suspension flow, J. Fluids Eng. 132, 011302 (2010).

[43] K. Yeo and M. R. Maxey, Numerical simulations of concentrated suspensions of monodisperse particles in a Poiseuille flow, J. Fluid Mech. 682, 491 (2011).

[44] U. Zimmermann, F. Smallenburg, and H. Löwen, Flow of colloidal solids and fluids through constrictions: Dynamical density functional theory versus simulation, J. Phys. Condens. Matter 28, 244003 (2016).

[45] M. R. Maxey, Simulation methods for particulate flows and concentrated suspensions, Annu. Rev. Fluid Mech. 49, 171 (2017).

[46] S. Mondal, C.-H. Wu, and M. M. Sharma, Coupled CFD–DEM simulation of hydrodynamic bridging at constrictions, Int. J. Multiphase Flow 84, 245 (2016).

[47] C. Bächer, L. Schrack, and S. Gekle, Clustering of microscopic particles in constricted blood flow, Phys. Rev. Fluids 2, 013102 (2017).

[48] F. Municchi, P. P. Nagrani, and I. C. Christov, A two-fluid model for numerical simulation of shear-dominated suspension flows, Int. J. Multiphase Flow 120, 103079 (2019).

[49] A. A. Herale, P. Pearce, and D. R. Hewitt, Emergent clogging of continuum particle suspensions in constricted channels, J. Fluid Mech. 1017, A24 (2025).

[50] A. Sauret, K. Somszor, E. Villermaux, and E. Dressaire, Growth of clogs in parallel microchannels, Phys. Rev. Fluids 3, 104301 (2018).

[51] G. Kelly and T. G. Fai, Multi-scale model of clogging in microfluidic devices with grid-like geometries, Proc. R. Soc. A: Math. Phys. Eng. Sci. 478, 20220119 (2022).

[52] A. R. Pries, T. W. Secomb, P. Gaehtgens, and J. F. Gross, Blood flow in microvascular networks: Experiments and simulation, Circ. Res. 67, 826 (1990).

[53] A. Erlich, P. Pearce, R. Plitman Mayo, O. E. Jensen, and I. L. Chernyavsky, Physical and geometric determinants of transport in feto-placental microvascular networks, Sci. Adv. 5, eaav6326 (2019).

[54] M. Šitina, H. Stark, and S. Schuster, Optimal hematocrit theory: a review, J. Appl. Physiol. 137, 494 (2024).

[55] V. Baranau and U. Tallarek, Random-close packing limits for monodisperse and polydisperse hard spheres, Soft Matter 10, 3826 (2014).

[56] H. H. Billett, Hemoglobin and hematocrit, in Clinical Methods: The History, Physical, and Laboratory Examinations, edited by H. K. Walker, W. D. Hall, and J. W. Hurst (Butterworths, Boston, 1990) 3rd ed.

[57] P. B. Canham and A. C. Burton, Distribution of size and shape in populations of normal human red cells, Circ. Res. 22, 405 (1968).

[58] P. Crucitti, V. Latora, and M. Marchiori, Model for cascading failures in complex networks, Phys. Rev. E 69, 045104 (2004).

[59] L. D. Valdez, Cascading failures in complex networks, J. Complex Netw. 8, cnaa013 (2020).

[60] D. Braess, A. Nagurney, and T. Wakolbinger, On a paradox of traffic planning, Transp. Sci. 39, 446 (2005).

[61] D. J. Case, Y. Liu, I. Z. Kiss, J.-R. Angilella, and A. E. Motter, Braess’s paradox and programmable behaviour in microfluidic networks, Nature 574, 647 (2019).

[62] A. Pinhas, J. V. Migacz, D. B. Zhou, M. V. C. Toral, O. Otero-Marquez, S. Israel, V. Sun, P. N. Gillette, N. Sredar, A. Dubra, J. Glassberg, R. B. Rosen, and T. Y. P. Chui, Insights into sickle cell disease through the retinal microvasculature: Adaptive optics scanning light ophthalmoscopy correlates of clinical oct angiography, Ophthalmol. Sci. 2, 100196 (2022).

[63] T. J. Brodribb, R. P. Skelton, S. A. M. McAdam, D. Bienaimé, C. J. Lucani, and P. Marmottant, Visual quantification of embolism reveals leaf vulnerability to hydraulic failure, New Phytol. 209, 1403 (2016).

[64] B. Schäfer, D. Witthaut, M. Timme, and V. Latora, Dynamically induced cascading failures in power grids, Nat. Commun. 9, 1975 (2018).

[65] H. M. Szafraniec, J. M. Valdez, E. Iffrig, W. A. Lam, J. M. Higgins, P. Pearce, and D. K. Wood, Feature tracking microfluidic analysis reveals differential roles of viscosity and friction in sickle cell blood, Lab Chip 22, 1565 (2022).

[66] H. M. Szafraniec, F. Bull, J. M. Higgins, H. A. Stone, T. Krüger, P. Pearce, and D. K. Wood, Suspension physics govern the multiscale dynamics of blood flow in sickle cell disease, Sci. Adv. 12, eadx3842 (2026).

[67] A. S. Baumgarten and K. Kamrin, A general fluid-sediment mixture model and constitutive theory validated in many flow regimes, J. Fluid Mech. 861, 721 (2019).

[68] F. Boyer, E. Guazzelli, and O. Pouliquen, Unifying suspension and granular rheology, Phys. Rev. Lett. 107, 188301 (2011).

