## Supplementary Material for "Clogging of particle suspensions in networks"

**This PDF file includes:**

- Supplementary figures
  - Supporting plots of solid flux and flow rate
  - Plots demonstrating robustness to the choice of $\phi_m$
    - Idealised network motifs
    - Complex network example
- Supplementary videos
  - Illustration of the network clogging algorithm

**Correspondence to:**
Duncan R. Hewitt
Philip Pearce

### 1 Supplementary figures

#### 1.1 Supporting plots of solid flux and flow rate ($\phi_m = 0.85$)

This subsection provides additional plots complementing Figs. 3 and 4 of the main manuscript. In Figs. S1 and S2, we plot the solid fluxes $\mathcal{F}_{ij}$ (panels (a)) and flow rates $Q_{ij}$ (panels (b)) in each vessel of the network, supplementing the corresponding plots of the ratio $\mathcal{F}_{ij}/Q_{ij}$ presented in the main manuscript.

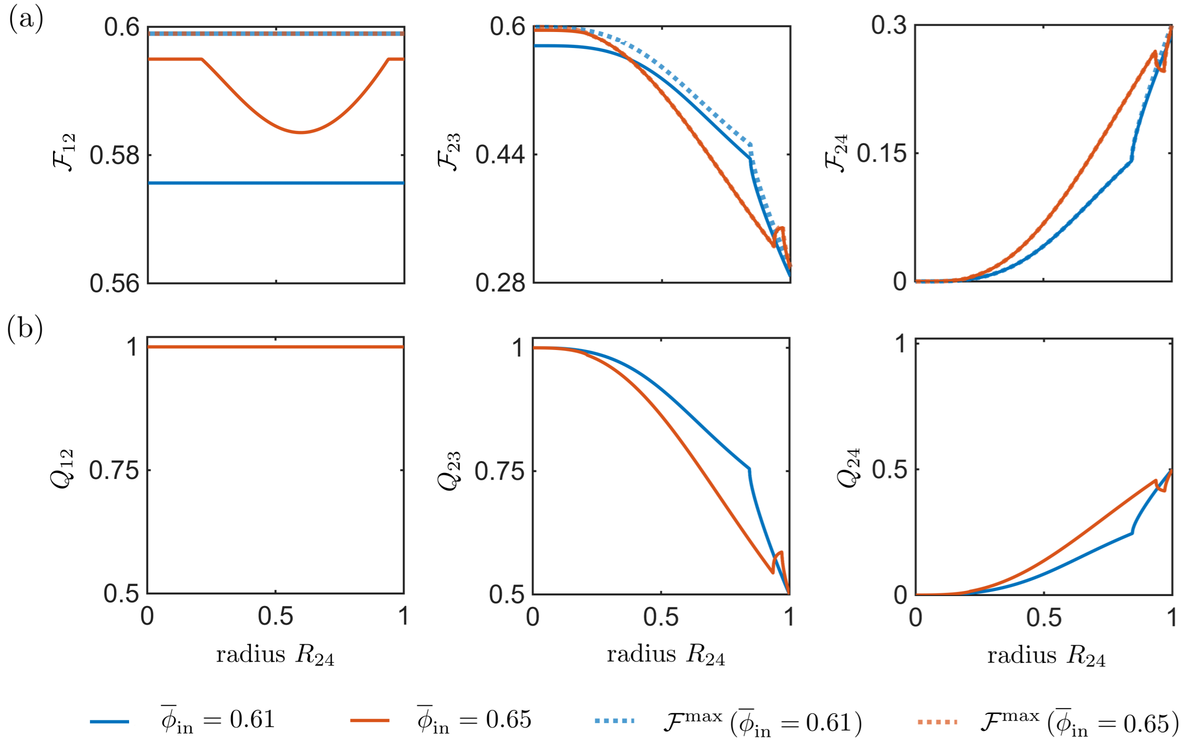


Fig. S1: Local redistribution mechanism in a three-vessel system with $Da = 10^{-3}$, $\phi_m = 0.85$ (complementing Fig. 3 of the main manuscript). (a) Solid fluxes $\mathcal{F}_{ij}$ (solid lines) and maximum solid fluxes $\mathcal{F}_{ij}^{\max}$ (dotted lines) in each vessel versus $R_{24}$. (b) Total flow rates $Q_{ij}$ in each vessel versus $R_{24}$. Panels (a–b) are shown for two values of $\bar{\phi}_{\text{in}}$.

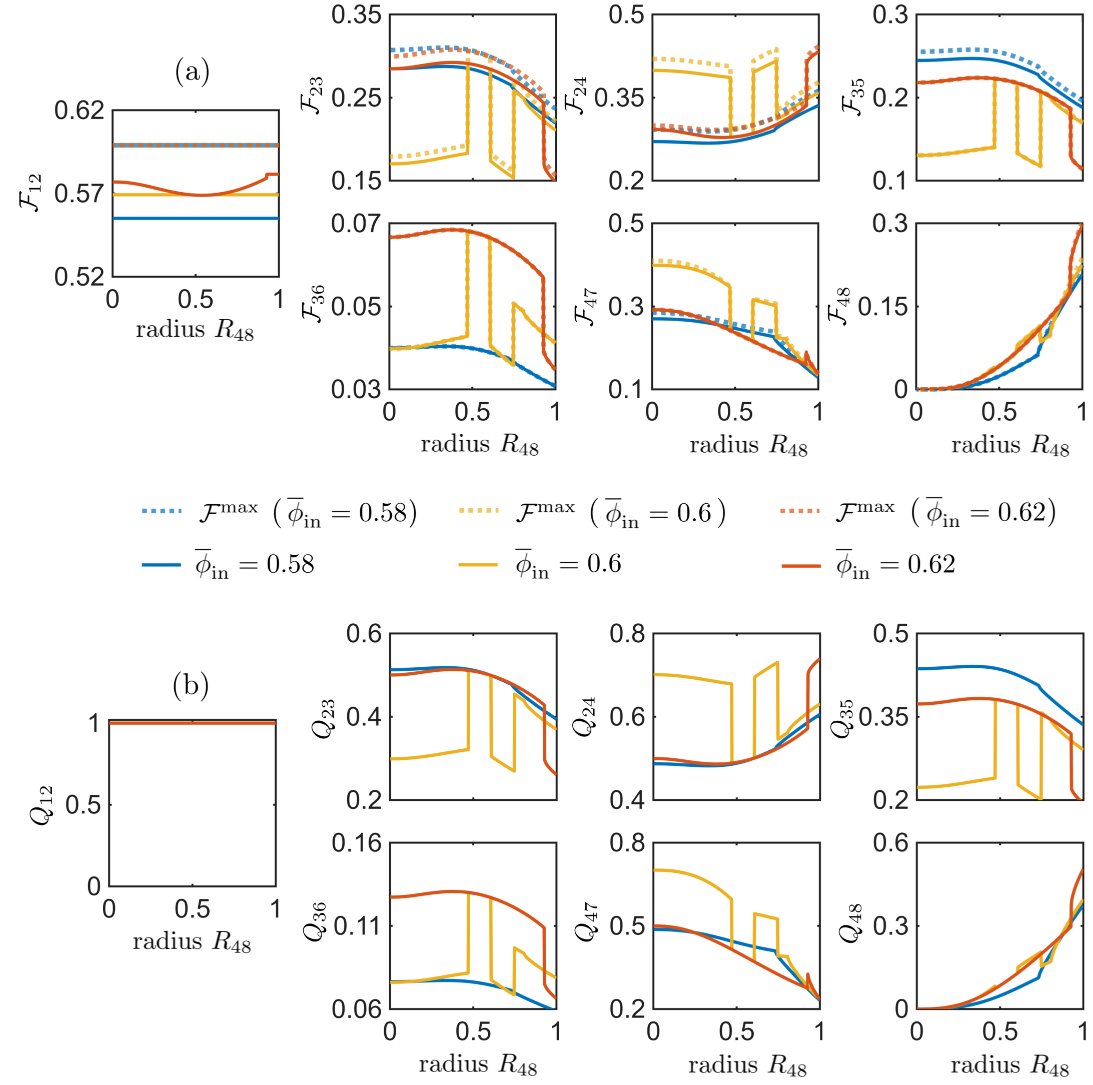


Fig. S2: Non-local clogging mechanism in a simple tree network with $Da = 10^{-3}$, $\phi_m = 0.85$ (complementing Fig. 4 of the main manuscript). (a) Solid fluxes $\mathcal{F}_{ij}$ (solid lines) and maximum solid fluxes $\mathcal{F}_{ij}^{\max}$ (dotted lines) in each vessel versus $R_{48}$. (b) Total flow rates $Q_{ij}$ in each vessel versus $R_{48}$. Panels (a–b) are shown for three values of $\bar{\phi}_{\text{in}}$.

#### 1.2 Plots demonstrating robustness to the choice of $\phi_m$

In the simulations presented in the main manuscript, the maximum packing fraction is $\phi_m = 0.85$, chosen as representative of deformable red blood cell suspensions. In this section, we demonstrate that the qualitative behaviour predicted by the model is preserved when using the alternative value $\phi_m = 0.65$, chosen to approximate the packing limit of rigid spherical particles. Here, we repeat the simulations from Figs. 3–6 of the main manuscript using this alternative value of $\phi_m$. The network radii are unchanged for Fig. 3, while slight adjustments are made for Figs. 4–5 to obtain comparable transport behaviour. In addition, for the complex network example (Fig. 6), we also consider the limiting case $\phi_m = 1$. While the quantitative thresholds for clogging are altered, the underlying mechanisms of local redistribution, non-local clogging, and topology-dependent transport remain unchanged, demonstrating that the conclusions of the main manuscript are robust to the choice of $\phi_m$.

##### 1.2.1 Idealised network motifs

In Figs. S3–S5, we reproduce Figs. 3–5 of the main manuscript for $\phi_m = 0.65$.

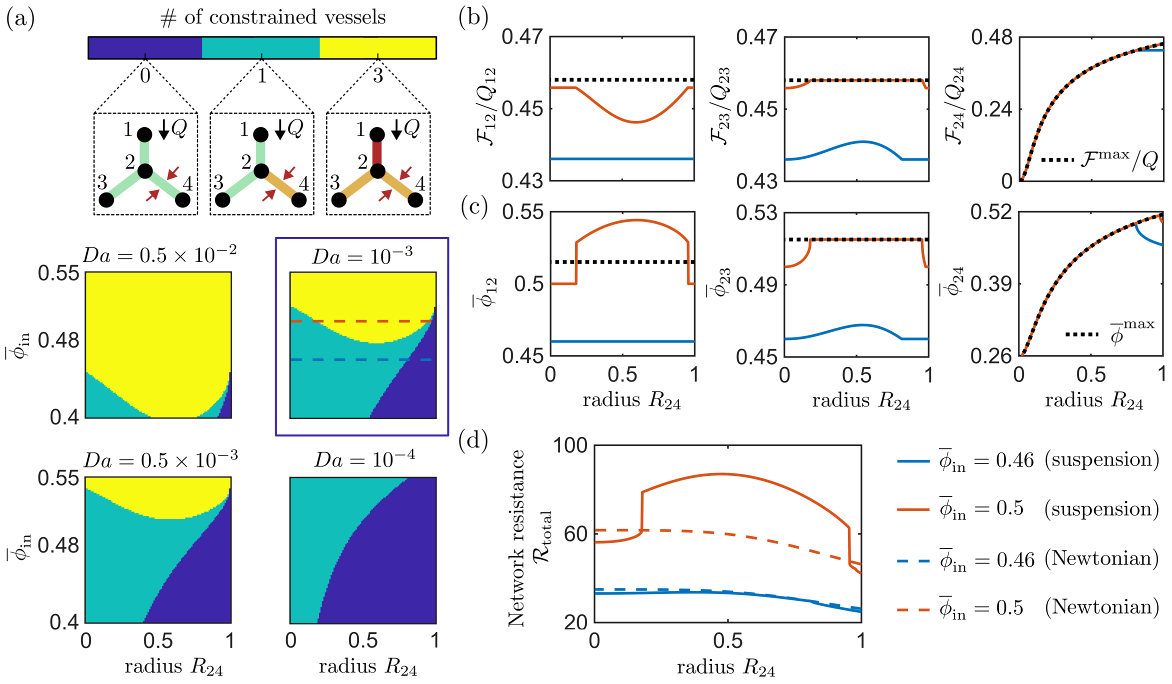


Fig. S3: Companion to Fig. 3 of the main manuscript, showing local redistribution (for the three-vessel network illustrated in panel (a)) for $\phi_m = 0.65$. The radius $R_{24}$ is varied while the remaining radii are held fixed at $R_{12} = 1$, $R_{23} = 1$. (a) Number of flux-constrained vessels as a function of $R_{24}$ and $\overline{\phi}_{\text{in}}$, shown for four values of $Da$. For $Da = 10^{-3}$, panels (b–d) show results for two values of $\overline{\phi}_{\text{in}}$ (indicated by dashed lines in (a)). (b) Solid fraction $\mathcal{F}/Q$ and $\mathcal{F}^{\max}/Q$ in each vessel versus $R_{24}$. (c) Particle volume fraction $\overline{\phi}$ and $\overline{\phi}^{\max}$ in each vessel versus $R_{24}$. (d) Total network resistance $\mathcal{R}_{\text{total}}$ versus $R_{24}$, with the corresponding Newtonian solutions shown with dashed lines.

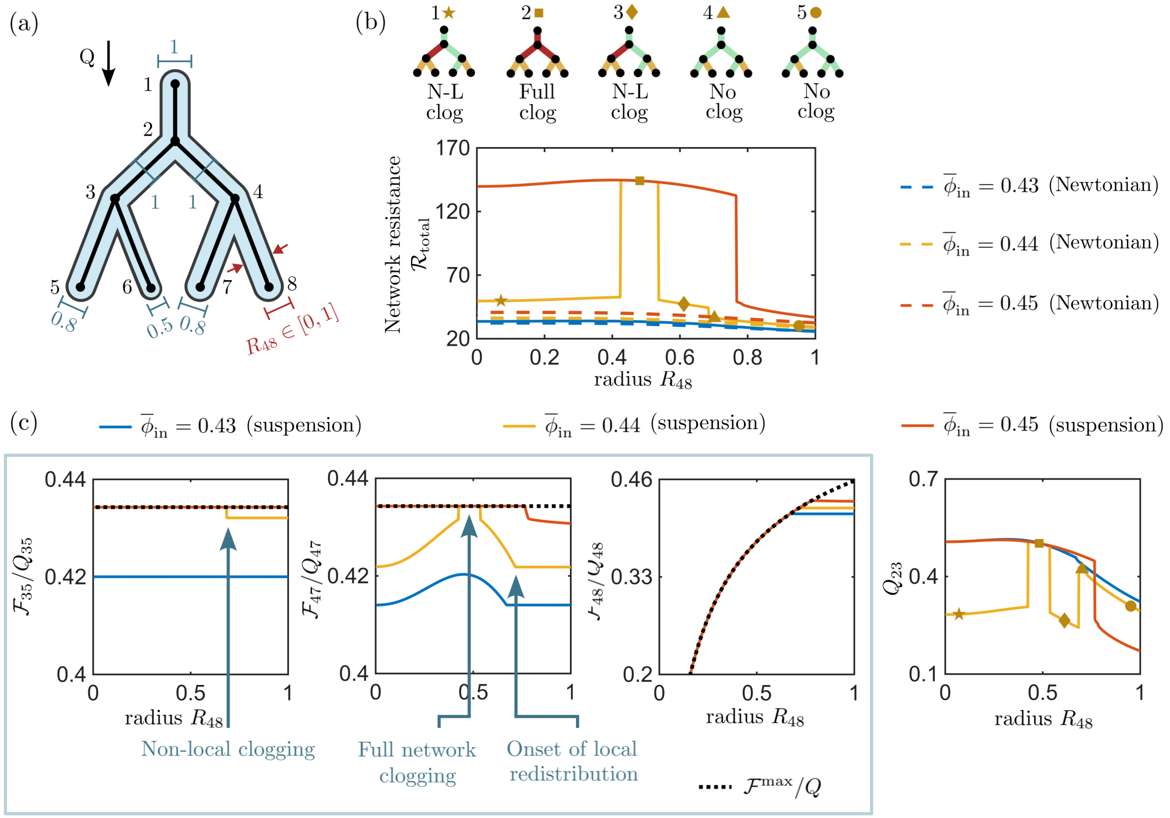


Fig. S4: Companion to Fig. 4 of the main manuscript, showing non-local clogging induced through variation in vessel radius in a simple tree network (see (a)) for $\phi_m = 0.65$. The network response is shown for three values of $\overline{\phi}_{\text{in}}$: a low value for which neither non-local or full network clogging occurs (blue), an intermediate value for which non-local clogging is followed by full network clogging (yellow), and a high value where the system transitions directly to full network clogging (red). Simulations are performed at $Da = 10^{-3}$. (b) Total network resistance $\mathcal{R}_{\text{total}}$ over $R_{48}$, with the corresponding Newtonian solutions shown by dashed lines. Network schematics 1–5 (left-right) indicate vessel states for the intermediate $\overline{\phi}_{\text{in}}$ case ($\overline{\phi}_{\text{in}} = 0.44$). (c) Solid fraction $\mathcal{F}/Q$ and $\mathcal{F}^{\max}/Q$ in three vessels ($\mathcal{V}_{35}$, $\mathcal{V}_{47}$, $\mathcal{V}_{48}$) versus $R_{48}$ (left), together with the total flow rate in vessel $\mathcal{V}_{23}$ (right).

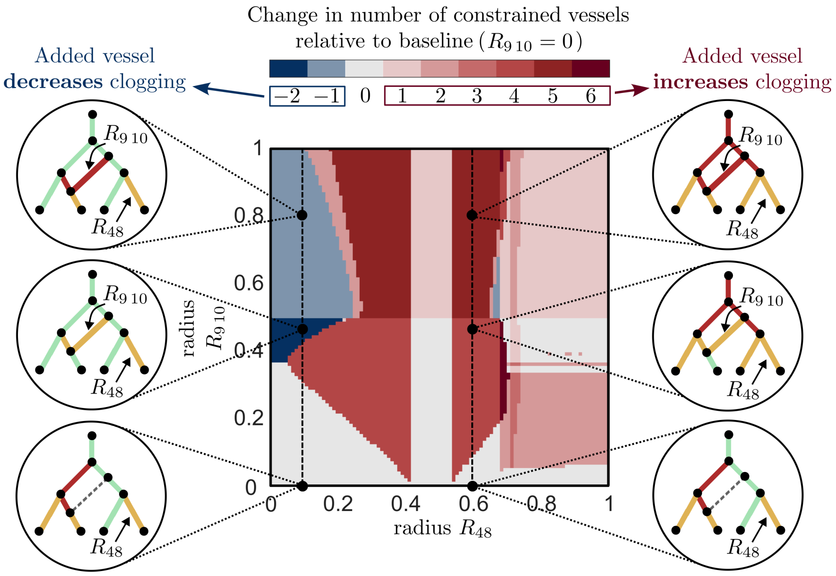


Fig. S5: Companion to Fig. 5 of the main manuscript, showing the effect of adding a bypass vessel for $\phi_m = 0.65$. Colormap shows the change in the number of constrained vessels as a function of vessel radii $R_{48}$ and $R_{9\,10}$, relative to the baseline case in which the bypass vessel is removed ($R_{9\,10} = 0$). Network geometry is identical to Fig. S4, with additional nodes 9, 10 introduced to split $\mathcal{V}_{24}$ and $\mathcal{V}_{36}$ respectively. Results are shown for $\overline{\phi}_{\text{in}} = 0.44$, $Da = 10^{-3}$, corresponding to the non-local clogging regime identified in Fig. S4 and also chosen to give a comparable degree of clogging to the $\phi_m = 0.85$ case shown in Fig. 5 of the main manuscript. The qualitative effect of introducing a bypass vessel on clogging is unchanged for this alternative value of $\phi_m$.

##### 1.2.2 Complex network example

In Figs. S6, we reproduce Fig. 6 of the main manuscript for the alternative maximum packing fractions of $\phi_m = 0.65$ and $\phi_m = 1$.

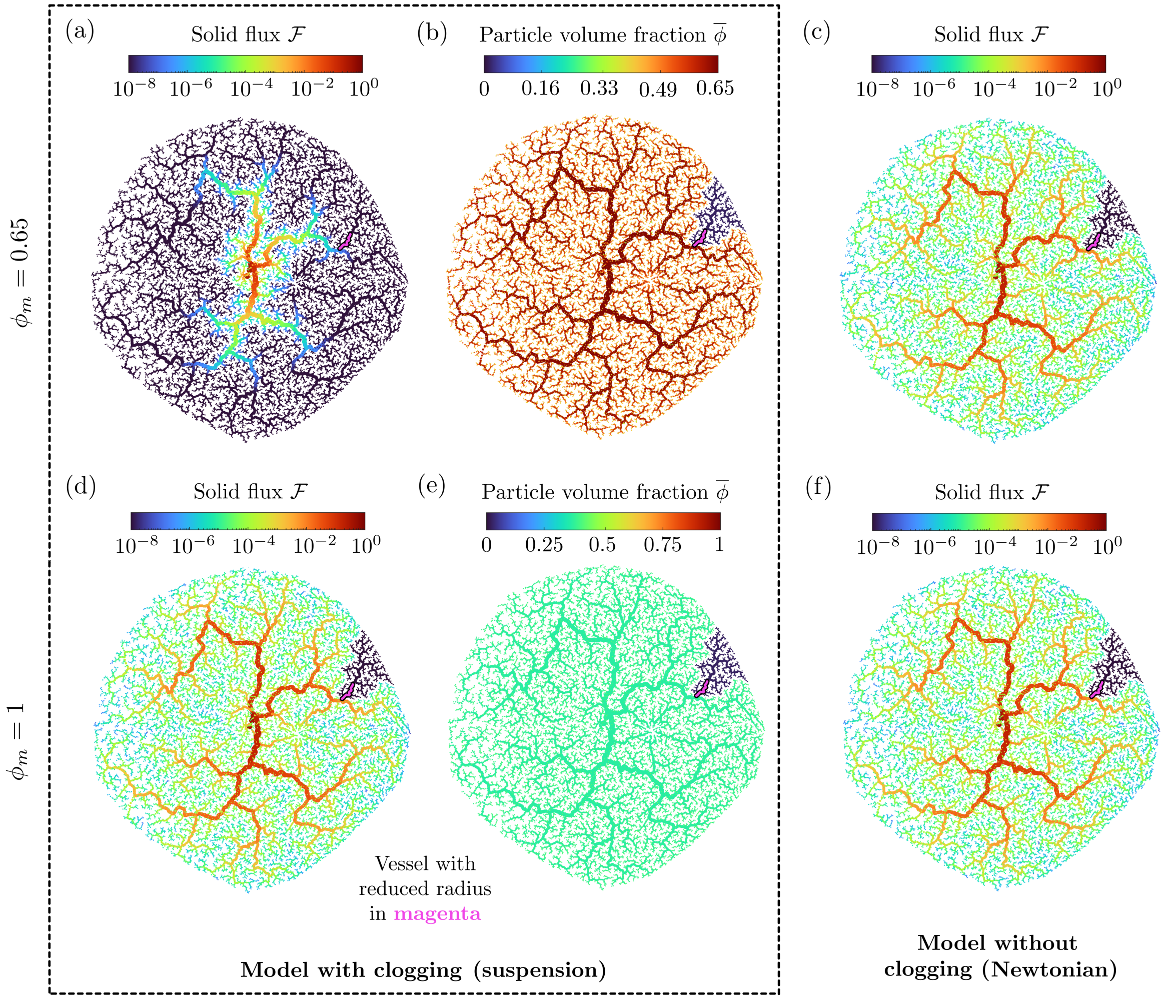


Fig. S6: Companion to Fig. 6 of the main manuscript, showing transport in the same complex network with the same short vessel segment (magenta) reduced to 1% of its original radius. Panels (a–c) show results for $\phi_m = 0.65$, while panels (d–f) show the corresponding results for $\phi_m = 1$. In both cases, $\overline{\phi}_{\text{in}} = 0.36$ and $Da = 10^{-5}$. Together with the intermediate $\phi_m = 0.85$ case shown in the main manuscript, these results demonstrate the influence of $\phi_m$ on clogging: decreasing $\phi_m$ promotes clogging, with most of the network clogged for $\phi_m = 0.65$, whereas no clogging occurs for $\phi_m = 1$. The choice $\phi_m = 0.85$ therefore provides an intermediate case in which clogging occurs without the entire network becoming clogged. (a,d) Solid flux distribution $\mathcal{F}$. (b,e) Particle volume fraction $\overline{\phi}$ for the suspension model. (c,f) Proxy for solid flux in a Newtonian fluid, defined as $\mathcal{F}_{ij} = \overline{\phi}_{\text{in}} Q_{ij}$.

### 2 Supplementary videos

#### 2.1 Illustration of the network clogging algorithm

**Video S1:** Iterative algorithm used to compute the clogged state of vessels in an example branching network (step 3 of Algorithm 1).

In Video S1, grey vessels indicate where the solid flux has not yet been calculated. During each iteration, solid fluxes are propagated downstream, with vessels operating below their maximum solid flux marked in green, and vessels set to their maximum solid flux marked in orange. Next, at each junction, if all downstream vessels are set to their maximum flux or are clogged, then all upstream vessels are set as clogged (marked in red). The solid flux is then recalculated in all vessels operating below their maximum solid flux, and the process repeats until no further vessels are set to their maximum flux, yielding the final clogged state.
